# miR-379-3p counteracts cancer cachexia through regulation of pyrimidinergic receptor, mitochondrial stress and interferon response

**DOI:** 10.1101/2025.01.11.632559

**Authors:** Maria Borja-Gonzalez, Raúl González-Ojeda, Anthony J Sannicandro, Chao Su, Elan C McCarthy, Clara Sanz-Nogués, Roisin M Dwyer, Brian McDonagh, Katarzyna Goljanek-Whysall

## Abstract

Cancer cachexia is a highly prevalent wasting syndrome in cancer patients. Inflammation is hallmarks of symptomatic cachexia, however early stages of cachexia are not well understood, including differences between biological sexes. In a mouse model of early cachexia, muscle from males showed strong mitochondrial defects, whereas females were characterized by inflammatory and stress response. We demonstrate a novel link between the increase in purinergic receptor P26Y, and dysregulated Ca^2+^ homeostasis, mitochondrial dysfunction and damage, and inflammation during early stages of cancer cachexia. Low levels of miR-379-3p were associated with poor survival of patients with lung cancer. Restoring miR-379-3p levels in mice prevented loss of muscle mass and function. miR-379-3p targeted P2r6y and restored mitochondrial content and function, inhibited type II interferon response, and regulated the expression of Ca^2+^-related and apoptotic markers. This supports miR-379-3p as a hub regulating multiple processes underlying cachexia and represent a therapeutic target for cancer patients.

## Introduction

Cancer cachexia (CC) is a severe catabolic syndrome characterized by loss of skeletal muscle [1, 2]. Despite the prevalence and severity, the mechanisms of CC are not well understood, and there are no effective therapies. Among potential therapeutic avenues explored at pre-clinical level are activation of AKT-mTORC1 signaling, blocking ActRIIB, NAD^+^ repletion and administration of mitochondrial antioxidant SkQ1 [3–6].

Muscle wasting in CC is associated with disrupted ubiquitin proteasome and lysosomal autophagy systems and decline in AKT activity with increased protein breakdown [7, 8]. Local and systemic inflammation, are considered hallmarks of CC [9, 10]. IL-6 and TNFα are implicated in the catabolic shift, particularly in female mice [11, 12]. Changes in mitochondrial homeostasis in CC have been demonstrated, occurring at earlier stages of cachexia in male than female mice [13–15]. The importance of biological sex on cancer cachexia is gaining recognition [3, 12, 13, 15–17] [18].

microRNAs (miRs), small non-coding RNAs that regulate multiple genes simultaneously and are regulators of muscle [19–21]. The levels of miRs change in muscle of cancer patients and in animal models [22–30]. Tumor-derived miRs have been proposed to mediate cachexia [31] and several studies proposed that regulation of miRs in CC may ameliorate muscle wasting [22, 28, 29, 32]. miR-379-3p is part of the DLK1-Dio3 cluster, and other miRs included in this cluster have been associated with tumor suppressor functions [33]. miR-379-3p is downregulated in *tibialis anterior* in a mouse model of cachexia (LLC) [30, 34], and in *vastus lateralis* of non-small cell lung cancer (NSCLC) patients [30, 35]. However, its physiological role remains poorly understood. The targets of miR-379-3p are largely unknown.

Here, we demonstrate the potential of miR-379-3p overexpression to ameliorate muscle loss in CC. miR-379-3p and its target: P2ry6, purinergic receptor associated with inflammation, were predictive of survival in lung cancer patients. Mice treated with miR-379-3p showed limited tumor growth, improved muscle size and function; these effects were more pronounced in females. The inflammatory and mitochondrial signature, and changes in apoptotic and Ca^2+^-related proteins in CC, demonstrated by proteomics and RNA-Seq analyses, were reversed by the treatment with miR-379-3p. Thus, restoring the levels of miR-379-3p has a therapeutic potential for ameliorating cachexia through coordinated regulation of purinergic signaling, mitochondrial damage and inflammatory response.

## Results

### Global proteomics reveal changes in energy metabolism in males and activation of type II interferon response in females in the early stages of cancer cachexia

Lewis Lung Carcinoma (LLC) mouse model was used to identify changes in CC. After three weeks of tumor growth (Fig.1A), female mice had larger tumors and greater, albeit limited, changes in body weight (BW) (Fig.1B). Muscle mass was reduced, with significant changes in quadriceps (QUAD) of cachectic female mice (LLC) compared to control (PBS) (Fig.1C). Higher proportion of smaller fibers was observed in LLC female mice compared to control, whereas grip strength (GS) was significantly decreased in LLC males (Fig.1C). This indicates the 3-week time point as onset of muscle loss in CC. In males, tumor growth was limited and only grip strength was significantly reduced in LLC compared to PBS mice (Fig.1B,C).

**Figure 1.**
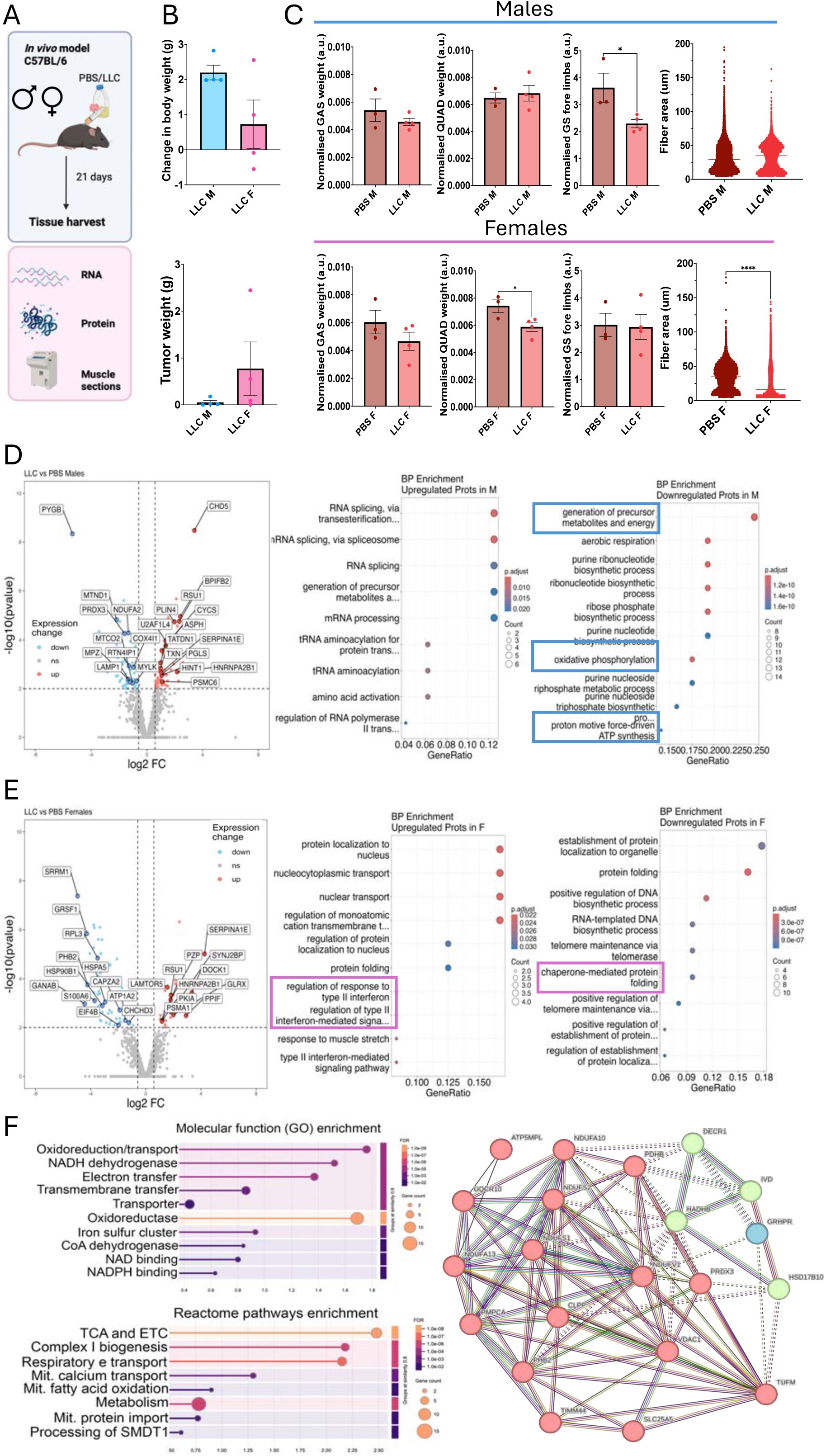
Phenotypic and proteomic changes in early stages of cancer cachexia. **A.** Study design of LLC model with subcutaneous injection of LLC cells or PBS in C57BL/6 mice. Tissue was harvested at day 21. **B.** Change in body weight (ΔBW) and tumour weight (TW) in males and females over 21 days (LLC n=4) was more pronounced in females. **C.** Normalised (to BW) muscle weights of gastrocnemius (GAS), quadriceps (QUAD) and grip strength (GS) at day 21 in PBS and LLC males and females (PBS n=3, LLC n=4). Minimal Feret’s in PBS and LLC males and females (PBS n = 3 and LLC n=4) show significant changes in fibre distribution in females. **D-E.** Changes in whole proteome in cachectic (LLC) mice compared to control (PBS) mice: volcano plots show significantly upregulated (red) and downregulated (blue) proteins. GO enrichment analysis for upregulated proteins in males and females highlights RNA translation, splicing, and inflammation pathways. GO enrichment analysis for downregulated proteins in males and females shows mitochondrial dysfunction and impaired protein folding. (PBS n = 4 and LLC n=4) **F. Up- or downregulated proteins common with MitoCarta 3.0 set indicate that a proportion of proteins regulated in mice during early stages of cachexia are associated with mitochondria.** Enrichment analysis (molecular function) shows proteins associated with oxidative phosphorylation electron transport chain and iron-sulfur clusters, and functional enrichment (Reactome) shows proteins regulated in CC were associated with TCA, complex I biogenesis, lipid metabolism and mitochondrial Ca^2+^ transport are regulated in males and females during CC. Data are shown as means with individual values. Error bars show SEM. Statistical analysis was performed using t-tests. *p<0.05; **** p<0.0001. A.U., arbitrary units. Individual data points are independent biological replicates (mice). PBS – control; LLC – cachectic mice; LLC miR-cachectic mice treated with miR-379-3p.

Global proteomics changes at the 3-week time point showed that upregulated proteins in males were associated with splicing, amino acid activation and tRNA (Fig.1D, Fig.S1A). Among these were: HNRNPA2B1, an RNA-binding protein regulating type I interferon responses (IFNα/β), such as IFI6 and cGAS, and CYCS and PSMC6, which control apoptosis. Downregulated proteins were associated with oxidative phosphorylation, ATP biosynthesis and respiration and included mitochondrial proteins, LAMP1 potentially indicating lysosomal defects, and PRDX3, associated with mitochondrial ROS (Fig.1D, Fig.S1A).

In LLC females, among upregulated proteins were SERPINA1E and PZP, associated with inflammation, SYNJ2BP associated with mitochondrial-ER membrane contact sites and PPIF, component of the mitochondrial permeability transition pore (Fig.1E). Downregulated proteins included heat shock proteins: HSP90AA1, HSP90AB1, HSP90AA1 and GRSF1, mitochondrial RNA-binding protein (Fig.1E). Upregulated proteins were enriched (GO term: biological function) for: type II IFN responses (IFNγ), whereas downregulated proteins were associated with regulation of DNA biosynthetic process, protein folding, telomerase, processing in the ER, necroptosis and pentose phosphate pathway (Fig.1E, FigS1B, D).

Pathways associated with mitochondria, inflammation, catabolism and structural muscle proteins, were enriched in muscle of both males and females during early stages of cachexia. In males only, splicing-associated genes were enriched in cachectic muscle, whereas in females only, there was a signature of ER stress in cachectic muscle.

Proteins regulated in males and females were compared with MitoCarta 3.0, composed of 1,141 known mitochondrial-associated genes [36]. This highlighted changes in mitochondria-associated proteins in both males and females (Fig.1F; Fig.S1D, Table S3). Enrichment analysis revealed that these proteins were associated with oxidative phosphorylation and electron transport chain, as well as iron-sulfur clusters, indicating changes in mitochondrial function (Fig.1F, Table S3). Functional enrichment indicated their association with TCA, complex I lipid metabolism and mitochondrial Ca^2+^ transport (Fig.1F, Table S3). Together, these data highlight the importance of mitochondrial changes during early stages of CC, with higher enrichment in males, and predominantly inflammation-related phenotype in females.

### High levels of miR-379-3p predict better outcomes in patients with lung cancer

We next cross-referenced cachexia-regulated proteins with microRNAs regulated in muscle during CC in humans and animal models [30]. miR-379-3p was downregulated in cachexia and its low expression was correlated with poor prognosis in lung squamous cell carcinoma (LUSC) patients (HR=0.47, 95% CI= 0.27 to 0.84, P= 0.0084), particularly in women with LUSC (HR=0.54, 95% CI= 0.29 to 1.01, P= 0.052) (Fig. 2A, Fig.S2). miR-379-3p was downregulated in QUAD from male and female tumor-bearing mice (Fig.2B). Overexpression of miR-379-3p in C2C12 myotubes co-cultured with LLC cells in a transwell system, to mimic CC *in vitro*, rescued decreased myotube diameter, area and fusion index (Fig.2C, D), but not myogenic differentiation (Fig.S3), suggesting that miR-379-3p may play an important role in CC through regulating muscle size.

**Figure 2.**
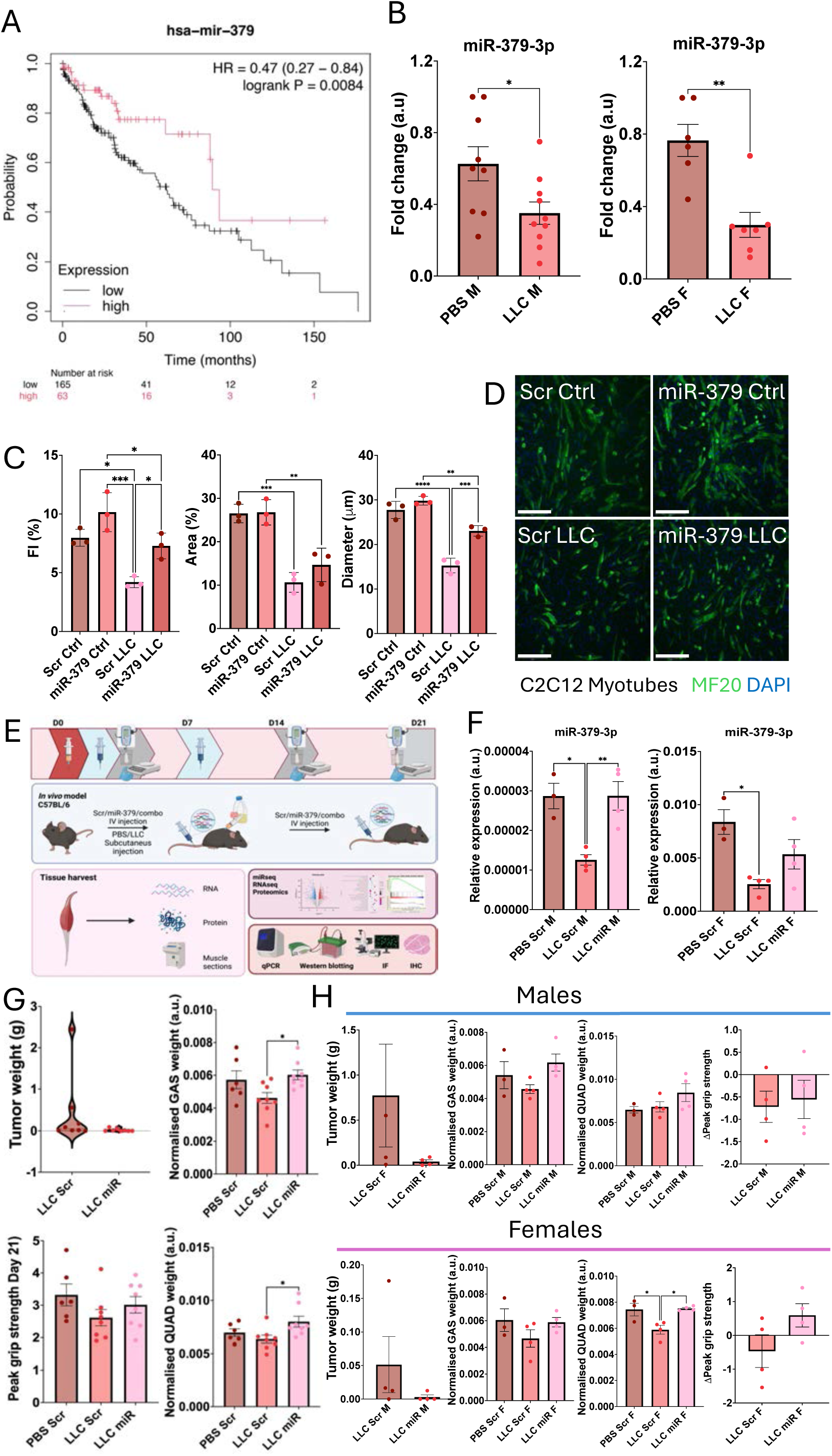
Restoring miR-379-3p levels during cachexia slows down cachexia progression in males and females. **A.** Survival curves generated from Kaplan-Meier plotter showed that low miR-379 expression was associated with worse prognosis in patients at the early stage of lung cancer (HR=0.47, 95% CI= 0.27 to 0.84, P= 0.0084). **B.** miR-379 expression was downregulated in QUAD from PBS and LLC male and female mice. n=9 of PBS M; n= 10 LLC M; n=6 of PBS F; n= 7 LLC F. **C-D.** miR-379 overexpression restores fusion index and myofiber diameter in *in vitro* model of cachexia: C2C12 myotubes co-cultured with LLC cells. MF20 immunostaining was performed, fusion index, FI, myotube area and the diameter of myotubes shown. Representative images are shown. N=3 replicates. Error bars show SD; * p < 0.05; ** p<0.01, *** p<0.001, **** p<0.0001; One-way ANOVA followed by Tukey’s multiple comparison test was performed to assess significances. Scale bar 275 μm. Analysis performed in ImageJ software. **E.** Study design of LLC model with subcutaneous injection of LLC cells or PBS in C57BL/6 mice. miR-379 injections were performed at D0 and D7. **F.** miR-379 expression was restored in LLC mice treated with miR-379-3p mimic delivered IV, compared to LLC mice. **G.** Restoring levels of miR-379-3p in cachexia slowed down tumor growth, preserved muscle mass and grips strength; data from male and female mice shown. **H.** miR-379-3p has more pronounced effects in females than males: tumor weight, normalised gastrocnemius (GAS), quadriceps (QUAD) in males and females shown in males and females separately. PBS n = 3, LLC n = 4, LLC miR n=4 for males and females each). qPCR: graphs showed relative expression to fold change (of aged-matched animals) or relative expression, normalised to Snord68 and Unisp6 average, analysis performed by ΔΔCT method. Error bars show SEM; *p<0.05 ** p < 0.01; Parametric, unpaired t-test and one-way ANOVA were used to analyse miR-379 expression. PBS – control; LLC – cachectic mice; LLC miR-cachectic mice treated with miR-379-3p.

### Restoring miR-379-3p levels during cachexia affects tumor growth and muscle mass in female mice

miR-379-3p levels were next restored in muscle of cachectic mice following miR-379-3p IV delivery (Fig.2E,F). Restoring miR-379-3p levels led to slower tumor growth and preserved muscle mass and partially grip strength (Fig.2G). Tumor size was reduced in both male and female LLC mice following miR-379-3p delivery and quadriceps mass was preserved in female LLC mice treated with miR-379-3p, together with grip strength (Fig.2H). These data indicate the potential for miR-379-3p to slow down cachexia progression, particularly in female mice.

### miR-379-3p regulates myofiber size and type in female mice with cancer cachexia

A higher proportion of the smallest fibers was detected in male and female mice cachectic mice (LLC); this was rescued by miR-379-3p in males (Fig.3A,B, Fig.S4A). A decrease in the proportion of middle-sized fibers in LLC mice was rescued by miR-379-3p in LLC females (Fig.3C, Fig.S4A). Fiber size distribution, but not the average diameter, was significantly changed in cachectic mice and rescued by miR-379-3p treatment (Fig.3B, Fig.S4). Analysis of myofiber types in TA muscle, mainly composed of fast types (IIa and IIb), affected during cachexia [1], showed decreased proportion of IIa fibers and increased proportion of type I fibers in female LLC mice; this was ameliorated by miR-379-3p (Fig.3D, Fig.S4C). In male mice, these changes in proportion of middle-sized fibres were not statistically significant (Fig.3D, Fig.S4C). H&E staining did not show obvious infiltration of immune cells in muscle of cachectic mice, however CD68 expression, but not CD206, was elevated, potentially suggesting presence of pro-inflammatory macrophages [37] and was significantly decreased in cachectic mice treated with miR-379-3p compared to LLC groups (Fig.3E,F; Fig.S4D,E).

**Figure 3.**
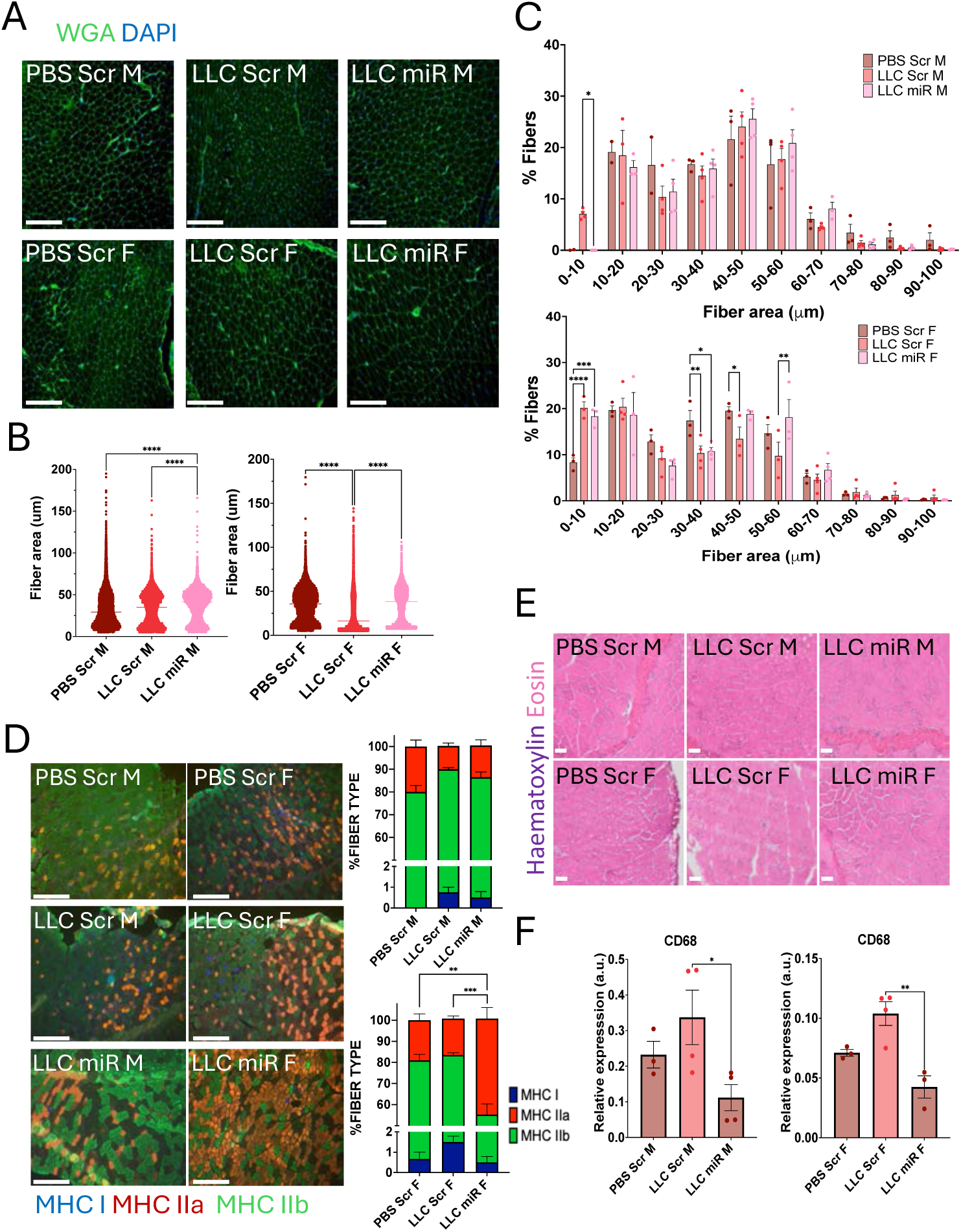
Restoring miR-379-3p levels during cachexia preserves myofiber size and predominantly in female mice. **A.** Representative images of WGA and DAPI immunostaining performed on GAS muscle shown. **B.** Fibre size and distribution represented per group in males and females show preserved fibre size distribution in cachectic males and females treated with miR-379-3p. **C.** Fibre percentage shows preserved middle range sized fibres, predominantly in cachectic females treated with miR-379-3p. **D.** miR-379-3p restores myofiber type proportion in cachectic females. TA cryosections immunostained using antibodies against MHC I (blue), MHC IIa (red) and MHC IIb (green). Representative images and percentage (%) of fibre type in males and females are shown. **E.** Haematoxylin and Eosin staining performed on TA does not reveal significant infiltration of immune cells. Representative pictures shown. **F.** Macrophage M1 marker, Cd68 was elevated in cachexia, but reduced in miR-379-3p treated male and female mice. qPCR on RNA isolated from QUAD; Cd68 relative expression was analysed using ΔΔCT method. Graphs showed relative expression to b2M (b2microgloulin). Error bars show SEM; PBS Scr, n=3; LLC Scr, n=4; LLC miR, n=4, for males and females each. Two-way ANOVA was used to assess differences among fibre size in each group and One-way ANOVA (Tukey’s test) for differences fibre size and among average fibre size. Chi Square test was performed for statistical difference in fiber type composition. Images were obtained using EVOS M7000 Imaging systems. Scale bar: 250 μm. H&E images: scale bar = 20 μm. One-way ANOVA followed by Tukey’s test was performed to assess significance in relative expression. *p<0.05; **p<0.01; *** p<0.001; ****p<0.0001. PBS – control; LLC – cachectic mice; LLC miR-cachectic mice treated with miR-379-3p.

### miR-379 regulates mitochondria-related proteins in males and type II IFN response in females

Global proteomics of QUAD revealed that in male LLC mice, miR-379-3p resorted some of the pathways dysregulated in CC (Fig. 1D, Fig.4A). Proteins upregulated following miR-379-3p treatment, were associated with oxidative phosphorylation and energy metabolism: MT-CO2, NDUFA2, NDUFA12, UQCRQ (Fig.4A, B). Downregulated proteins in miR-379-3p-treated LLC males, compared to LLC males, were involved in glycolysis/gluconeogenesis, suggesting a downregulation of glucose metabolism (Fig.4A, B). MYH7, associated with type I fibers, was reduced after miR-379-3p treatment, consistent with earlier findings of type I fiber downregulation in LLC miR mice (Fig.2A, 4A).

**Figure 4.**
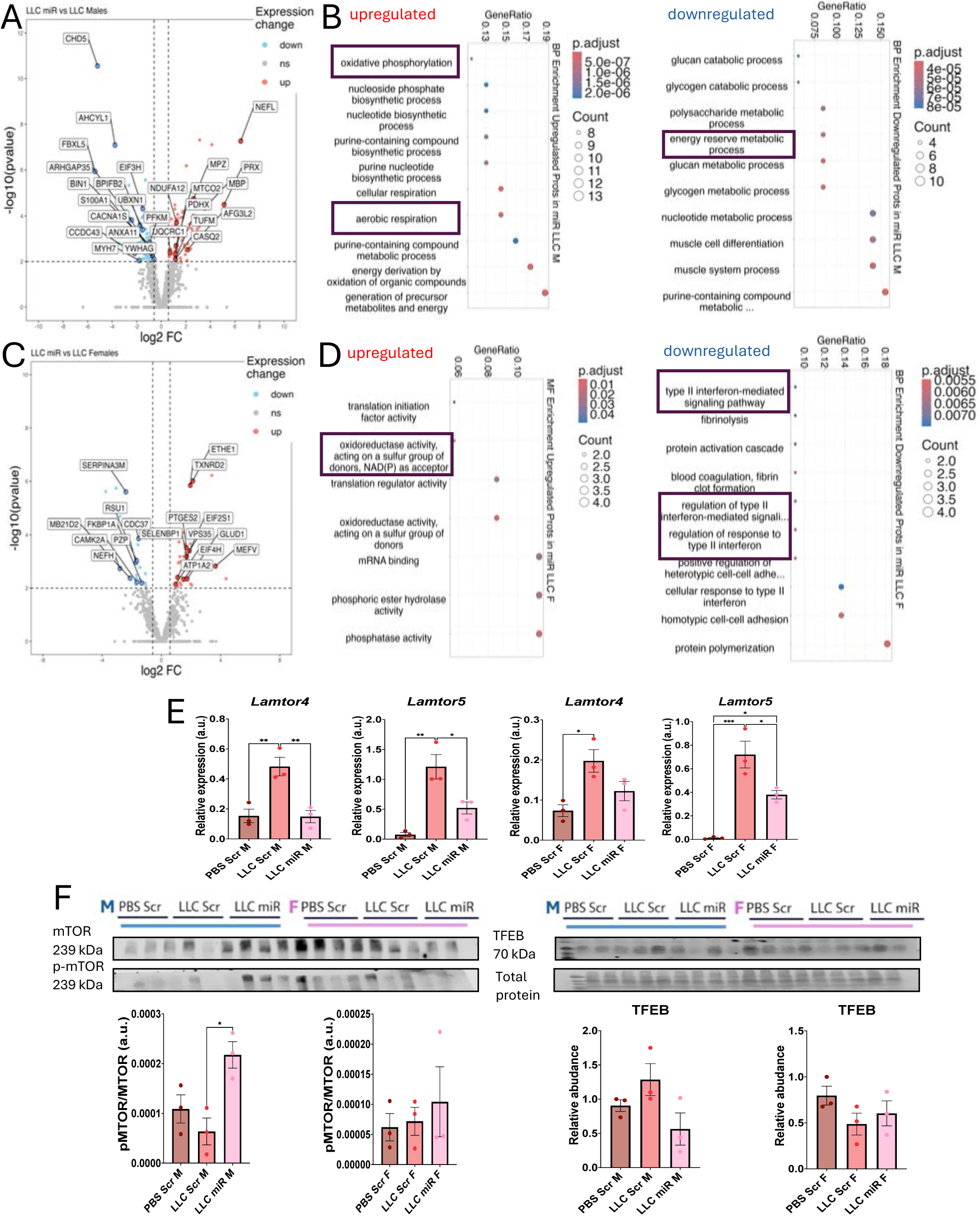
Global proteomics reveals miR-379-3p as a master regulator of mitochondria, inflammation and P2ry6 expression in muscle during cancer cachexia. **A., C.** Volcano plots showing differentially abundant proteins in the muscle of LLC miR-treated male (A) and female (C) mice. **B., D.** GO term enrichment analyses show that proteins regulated by miR-379-3p in male (B) cachectic mice are associated with energy metabolism and mitochondria, whereas in cachectic females (D), miR-379-3p regulated proteins were associated with type II interferon response and metabolism (see also Figure 2). N = 4. **E.** Nutrient sensing pathway is altered in cachexia and partially restored by miR-379-3p. The expression of Lamtor4, Lamtor5 (regulating assembly of mTOR complex), was upregulated in cachexia and partially restored by miR-379-3p treatment, Expression relative to b2M (b2microgloulin) analysed using ΔΔCT shown. N = 3 **F.** Phosphorylated mTOR protein was decreased in muscle of cachectic male and increased in miR-379-3p cachectic male, but not female mice. The levels of TFEB lysosomal biogenesis factor, affected by mTOR activity, showed increasing trend in cachectic male mice but not female that was restored by miR-379-3p delivery. Western Blots shown (n=3 in each group), abundance normalised to all protein (Ponceau S). Bar graphs showed densitometry analysis, performed in ImageJ software. Error bars showed SEM. One way ANOVA was performed. * p < 0.05; ** p< 0.01; *** p<0.001; ****p<0.0001. PBS – control; LLC – cachectic mice; LLC miR-cachectic mice treated with miR-379-3p.

In the muscle of female LLC miR-treated mice, downregulated proteins included FGA, FGB, CDC37, PZP, CDC151, and NEFH, associated with IFN-γ response, fibrinolysis, and wound healing; MB21D2, enhancing cGAS activity [38] and CAMK2A, promoting mitochondrial Ca²⁺ influx [39] (Fig.4B). Proteins upregulated in LLC miR-treated females included PTEGS2, SELEPB1, and TXNRD2, associated with oxidoreductase activity; EIF2/4, involved in peptide metabolism; and GLUD1, regulating energy homeostasis (Fig.4B). Moreover, PPIF, a part of the mitochondrial permeability transition pore (mPTP) regulating Ca^2+^ flux, GLRX, and EIF4B were altered, with chaperone-mediated folding and interferon response regulation pathways enriched (Fig. S5A). This suggests the role of miR-379-3p in protecting muscle during CC through the regulation of mitochondria and inflammation.

Energy and metabolism-associated pathways were regulated by miR-379-3p in muscle from both male and female mice and their disruption is a hallmark of symptomatic cachexia. The expression of Lamtor4 and Lamtor5, important for Ragulator assembly, mTOR signaling and innate immune response [40], was upregulated in muscle of cachectic males and females and downregulated following miR-379-3p treatment in LLC mice (Fig.4E). mTOR phosphorylation was decreased in LLC male, but not female, mice and was restored by miR-379-3p (Fig.4H). TFEB levels were not significantly affected, but showed trends consistent with changes in phosphorylated mTOR (Fig.4H).

### miR-379-3p regulates mitochondrial content and function in cancer cachexia

Based on the strong mitochondrial signature, proteins regulated by miR-379-3p in cachectic mice were cross-referenced with MitoCarta3.0 (Table S4). miR-379-3p regulated mitochondrial protein associated with metabolism and ETC in males (Fig.5A) and Krebs cycle and redox homeostasis in females (Fig.5B). The levels of Nd1, a mitochondria-encoded gene, and TOM20, a nuclear-encoded mitochondrial protein, were decreased in muscle of cachectic mice and partially restored by miR-379-3p treatment (Fig. 5C, D), suggesting prevention of mitochondrial loss by miR-379-3p in cachexia (Fig.5C, D).

**Figure 5.**
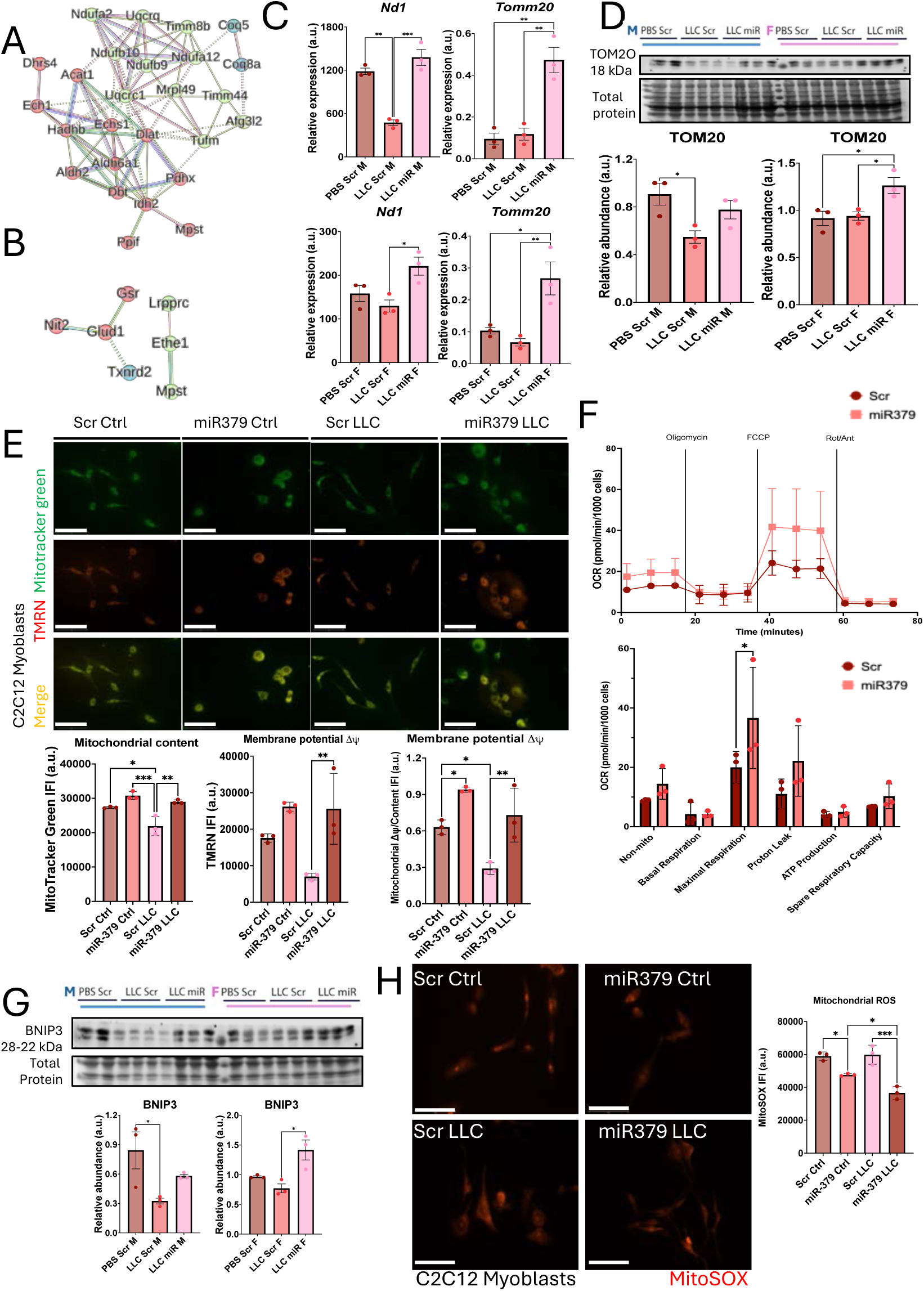
miR-379 regulates mitochondrial homeostasis in muscle during cancer cachexia. **A.,B.** STRING PPI network was represented of mitochondrial-associated proteins increased in LLC miR-379-3p treated males (A) and females (B). **C., D.** Mitochondrial content markers: Nd1 and Tomm20 are reduced in muscle of cachectic mice and restored by miR-379-3p treatment. Expression relative to b2M (b2microgloulin) shown. Western blot (n=3 for each group) graphs used for densitometry analysis, performed in ImageJ. **E., F.** miR-379-3p regulates mitochondrial content and function in *in vitro* model of cachexia. **E.** C2C12 myoblasts were co-cultured LLC cells or with DM (Ctrl), and treatment of Scr/miR-379 mimics was performed at the same time. MitoTracker green and TMRM staining representative images shown. Integrated fluorescence intensity of Mitotracker green, TMRM probe and analysis of the ratio of both parameters were performed in ImageJ. **F.** Seahorse bioanalysis of oxygen consumption of myoblasts was assessed, showing increased OCR in miR-379 treated C2C12s myoblasts. OCR parameters were assessed independently. N=3. **G.** Western Blot and densitometry analysis, performed in ImageJ show significant changes in mitophagy marker: BNIP3 in muscle of cachectic and cachectic mice treated with miR-379-3p. PBS n=3; LLC and LLC miR n=3. **H.** MitoSOX staining shows reduced mtROS production in C2C12 cells treated with miR-379-3p, with or without LLC co-culture. N=3. Representative images shown. Graphs showed densitometry analysis, performed in ImageJ. Scale bar 75 μm (E) and 50 μm (H). Error bars showed SEM (qPCRs and WB) and SD (cell probes and Seahorse); * p < 0.05; ** p < 0.01; *** p<0.001 One-way ANOVA followed by Tukey’s multiple comparison test and Kruskal-Wallis’s test were conducted to determine significance. Mann-Whitney test was used to assess the significance in OCR. Two-way ANOVA was performed to assess the significance in OCR parameters after normality was checked. PBS – control; LLC – cachectic mice; LLC miR-cachectic mice treated with miR-379-3p.

In an *in vitro* cachexia model, mitochondrial content, assessed by Mitotracker Green, was reduced in myoblasts co-cultured with LLC cells, and restored by miR-379-3p overexpression (Fig.5E). Mitochondrial membrane potential, measured by TMRN assay, normalized to mitochondrial content, was also reduced in C2C12 cells co-cultured with LLC cells, and rescued by miR-379-3p overexpression (Fig.5E).

Seahorse analysis validated miR-379-3p positive regulation of mitochondrial function: a significant increase in oxygen consumption rate (OCR) and basal and maximal respiration were observed in miR-379-3p-treated cells as compared to control (Scr) (Fig.5F), supporting the increase in mitochondrial content and function.

To determine whether changes in mitochondrial content and function may be related to changes in mitochondrial turnover, the levels of proteins associated with autophagy (Lc3bII/I, p62), biogenesis (Opa1) and fusion (Mfn2) were analyzed. These were not significantly affected by cancer cachexia or miR-379-3p (Fig.S5C). However, the levels of BNIP3, associated with mitophagy and apoptosis [41] were downregulated in LLC mice and restored to basal levels in mice treated with miR-379-3p, compared to LLC mice (Fig.5G), suggesting limited regulation by miR-379-3p at early stages of cachexia. This is consistent with previous finding showing increase in BNIP3 at later stages of cachexia [13]. Finally, mtROS production estimated with MitoSOX demonstrated a decrease in mtROS following miR-379 treatment in Ctrl and C2C12 co-cultured with LLCs, compared to Scr-treated C2C12 myoblasts (Fig.5H). Together, these data demonstrate that mitochondrial homeostasis is dysregulated in muscle of male and female cachectic mice, with male mice showing a stronger mitochondrial phenotype, and potentially different mechanisms associated with mitochondrial changes in female mice.

### miR-379-3p regulates inflammation in cancer cachexia

In male cachectic mice, mitochondria-related genes were predominantly regulated and restored by miR-379-3p (Figs.4,5). These changes were less pronounced in female mice. To further understand the cachexia phenotype in female mice, we performed RNA-Seq (Fig.6). Gene set enrichment analysis (GSEA) indicated upregulation of oxidative phosphorylation and mTORC1 signaling pathways, critical for ATP production and mitochondrial biogenesis (Fig.S5D). Moreover, inflammatory pathways related to Toll-like receptor (TLR) and TNFα signaling were dysregulated in female cachectic mice, whereas miR-379-3p treatment resulted restored the expression of genes associated with these pathways (Fig.6A,B,C).

**Figure 6.**
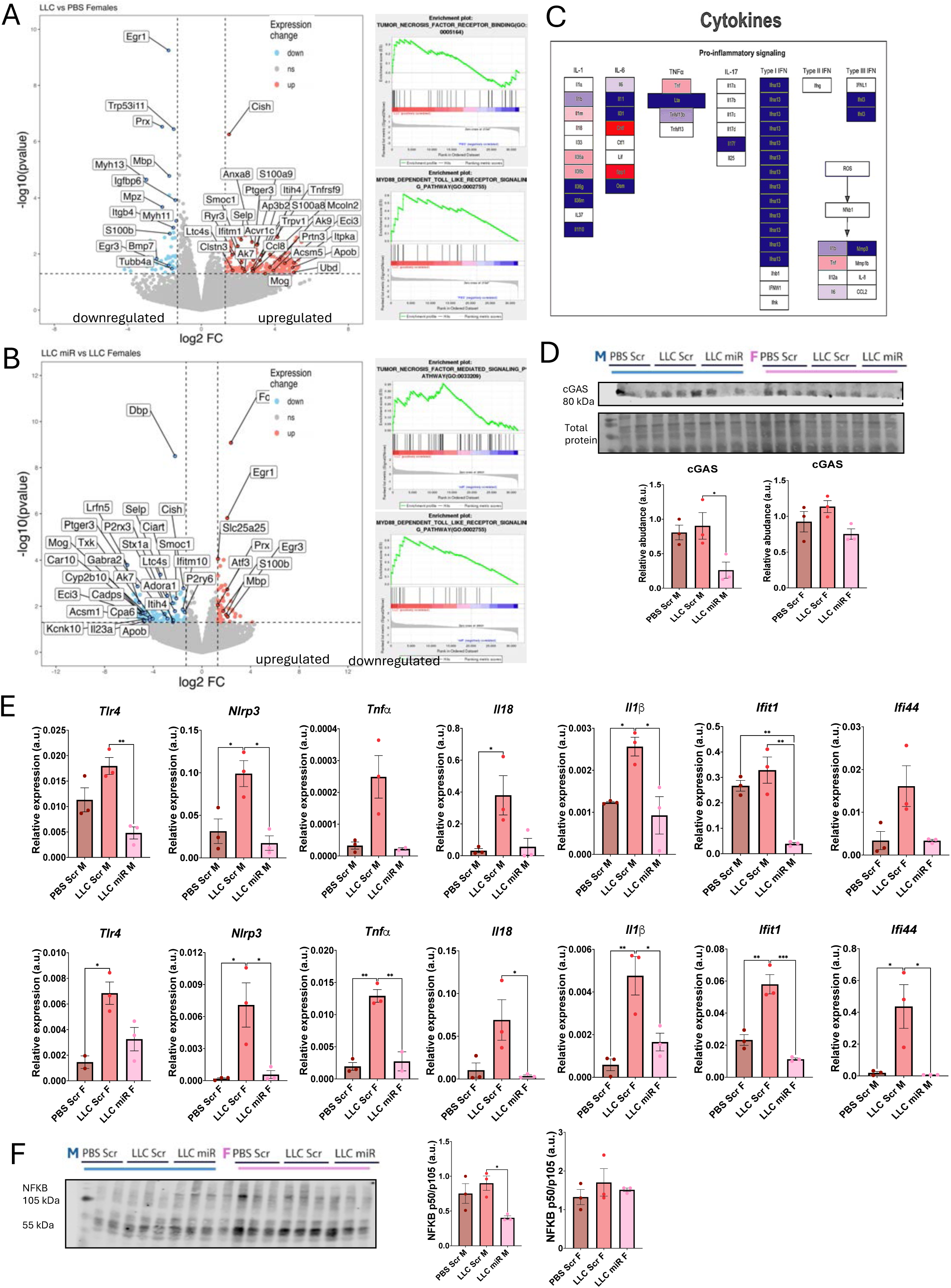
miR-379 modulates inflammation-associated pathways during cancer cachexia in females. **A.** RNA-seq on QUAD from LLC vs PBS female mice. Volcano plot shows differentially expressed genes in muscle from LLC vs PBS. Upregulated (red) genes reveal regulation of inflammation and immune pathways. Gene set enrichment analysis (GSEA) highlights significant pathways including tumour necrosis factor (TNF) signalling and MyD68 receptor signalling as enriched in the muscle of LLC female mice. **B.** Volcano plot shows differentially regulated genes in muscle from LLC miR vs LLC female mice. Downregulated genes (blue) reveal regulation of inflammation and immune pathway. Gene set enrichment analysis (GSEA) revealed tumour necrosis factor (TNF) signalling, MyD68 receptor signalling and as negatively enriched in the muscle of LLC miR female mice, compared to LLC mice. **C.** PathVisio pathway analysis of DEGs revealed the downregulation of genes involved in inflammation after miR-379 treatment in LLC female mice. Blue-downregulated, red – upregulated mRNAs after miR-379-3p treatment of cachectic mice. LLC n=4, PBS n=4. **D.** Western blot shows upregulation in cGAS (associated with DAMPs) levels in muscle of cachectic mice, which was reduced in the muscle of miR-379-3p treated cachectic male, but not female mice. **E.** Inflammation-related genes (*Tnfα, Il-18* and *Tlr4)* and inflammasome-related genes (*Nlrp3, Il-1β, Ifit1* and *Ifi44)* relative expression to *b*2M (*b*2microgloulin) housekeeper measured using ΔΔCT method in males and females. **F.** The levels of NF-kB, master regulator of inflammation, were upregulated in muscle during cachexia and restored by miR-379-3p treatment in male mice; NF-kB levels show similar trends in female mice. N=3. N=3, Error bars showed SEM (qPCRs and WB); * p < 0.05; ** p < 0.01; *** p<0.001 One-way ANOVA followed by Tukey’s multiple comparison test. PBS – control; LLC – cachectic mice; LLC miR-cachectic mice treated with miR-379-3p.

In cancer cachexia, inflammatory response has been shown to be downstream of TLR4 activation associated with the release of danger-associated molecular patterns (DAMPs), including mitochondrial DNA (mtDNA) [42]. We investigated changes in cGAS levels, a protein which functions as a sensor of double-stranded DNA fragments, including cytosolic mtDNA, and initiates an immune response *via* the adaptor protein STING. The levels of cGAS were elevated in cachectic male mice (LLC), whereas miR-379-3p treatment reduced the levels of cGAS (Fig.6D). A similar trend was observed in female mice (Fig.6D). This suggests mitochondrial damage, and a link to inflammation.

Based on RNA-Seq data suggesting activation of inflammatory pathways, and cGAS upregulation, we analysed the expression of Tnfa, Il1b and Il18 – pro-inflammatory cytokines; Ifit1and Ifi44 – interferon-regulated genes, and Tlr4 and Nlrp3, genes related to inflammasome. These genes were increased in muscle of cachectic female and surprisingly male mice and the treatment with miR-379-3p reversed these cachexia-induced changes (Fig.6E). Consistently, The levels of NF-kB (p50/p110 ratio), a key regulator of inflammation in muscle, were elevated in muscle from cachectic males, but not significantly in females, and reduced following the treatment with miR-379-3p (Fig.6F). This supports an important role of miR-379 in regulating inflammation during cachexia.

### miR-379-3p regulates the expression of apoptosis-related genes in cachectic mice

In addition to inflammatory changes, RNA-seq data analysis revealed a downregulation of apoptosis-related genes in miR-379-3p-treated cachectic female mice (Fig.7A). Moreover, upregulation of cGAS (Fig.6D) suggests mitochondrial damage which activates Caspase cleavage and apoptosis. Consistently, the expression of Bax, a pro-apoptotic regulator, was elevated in LLC mice but significantly reduced in LLC miR-treated mice, and Bak expression showed a similar trend (Fig.7B). The expression of Bcl2, an anti-apoptotic factor, was downregulated in cachectic mice and restored by miR-379-3p treatment (Fig.7B).

**Figure 7.**
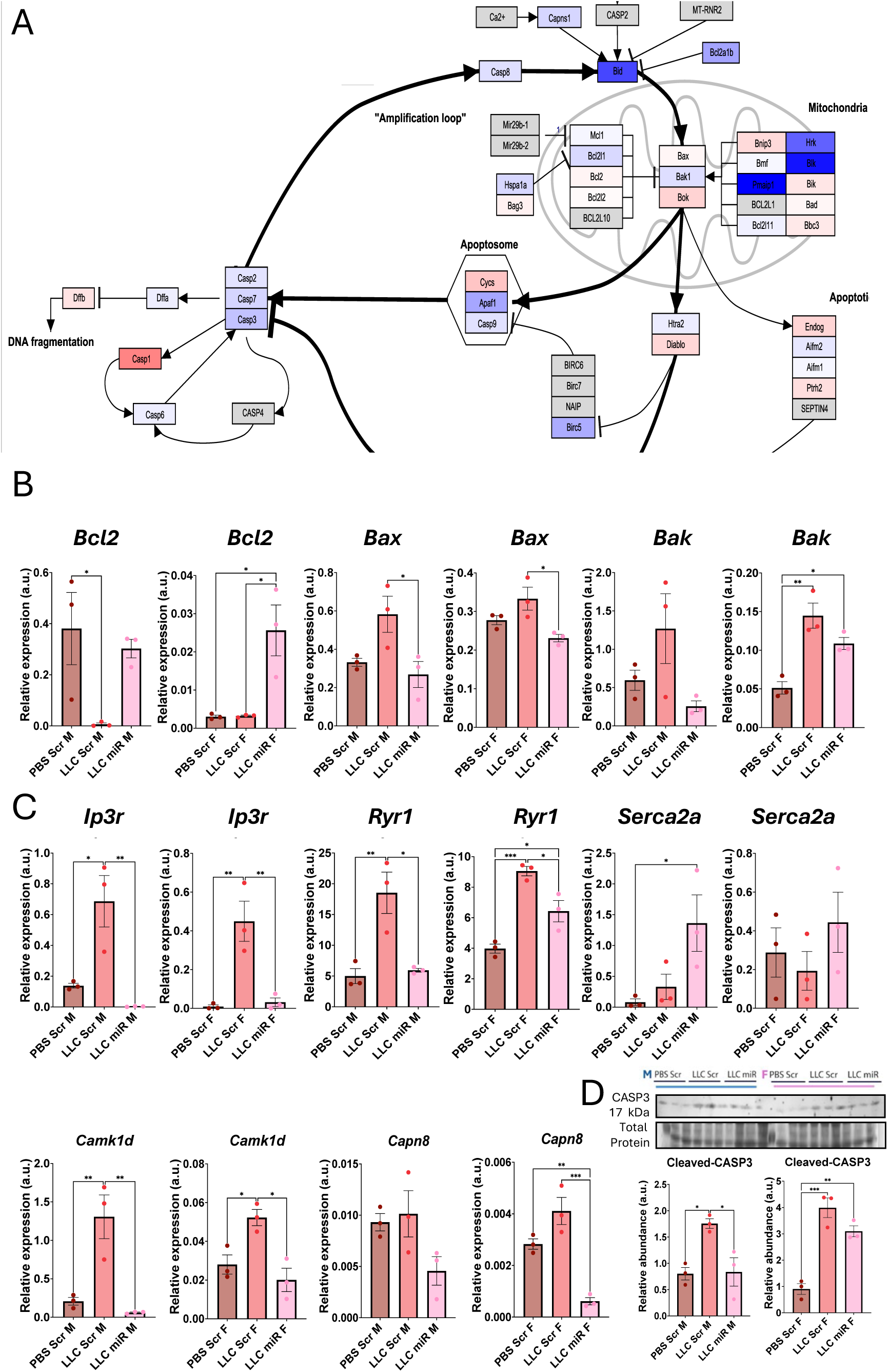
miR-379 regulates genes associated with apoptosis and calcium signalling during cancer cachexia. **A.** PathVisio pathway analysis of DEGs revealed a downregulation of genes involved in apoptosis signalling. Genes with a p-value cut-off of 0.05, represented as -log10 p-value, log 2-fold change cut-off ± 1 were included in Pathvisio software. **B.** Intrinsic apoptotic pathway related genes (*Bcl2, Bax* and *Bak)* expression relative to b2M (b2microgloulin) show upregulation of pro-apoptotic factors during cachexia in muscle of male and female mice, this was ameliorated by miR-379-3p treatment. **C.** Calcium signalling regulators related genes (*Ip3r, Ryr1 Serca2a, Camk1d* and *Capn8*) expression relative to b2M (b2microgloulin) shows upregulation of Ip3r and Ryr in muscle of male and female cachectic mice, suggesting disruption of flow of Ca^2+^ between ER and mitochondria, potentially activating apoptotic pathways. This was ameliorated by miR-379-3p delivery in both male and female cachectic mice. **D.** Western blot analysis shows increased cleavage of CASPASE-3 indicating increased pro-apoptotic signalling. One-way ANOVA followed by Tukey’s multiple comparison test and Kruskal-Wallis were performed to assess significance. PBS Scr, n=3 males and females; LLC Scr, n=3 males and females and LLC miRs, n=3 for males and females. PBS – control; LLC – cachectic mice; LLC miR-cachectic mice treated with miR-379-3p.

Proteomics analyses revealed changes in PPIF, a protein forming a part of the mitochondrial permeability transition pore (mPTP). Opening of the mitochondria may result from disrupted Ca^2+^ flow, which can trigger the opening of mPTP, activating pro-apoptotic pathways. RNA-Seq analyses showed changes in several genes related to Ca^2+^ homeostasis in female cachectic mice (Fig.6A,BC). The levels of Ip3r, encoding for inositol trisphosphate receptor, a membrane glycoprotein complex acting as a Ca^2+^ channel, and Ryr1, encoding for ryanodine receptor 1 regulating Ca^2+^ release, were upregulated in cachexia in male and female mice, and this upregulation was ameliorated by miR-379-3p (Fig.7C). Serca2 levels, encoding for sarcoplasmic/endoplasmic reticulum Ca^2+^-ATPase, were upregulated in cachectic mice treated with miR-379-3p, particularly in male mice (Fig.7C). Camk1d, encoding for calcium/calmodulin-dependent protein kinase 1 and Capn8, regulating Ca^2+^-dependent cysteine-type endopeptidase activity, were also upregulated in muscle from cachectic male and female mice, and this was rescued by miR-379-3p overexpression (Fig.7C). Finally, Western Blot analysis revealed an increase in cleaved CASP3, an indicator of apoptosis [43], in cachectic male and female mice, whereas miR-379 treatment significantly reduced CASP3 activation (Fig.8D), further cementing the critical role of miR-379-3p in regulation of Ca^2+^ homeostasis, mitochondria, inflammation and apoptosis, during cancer cachexia.

**Figure 8.**
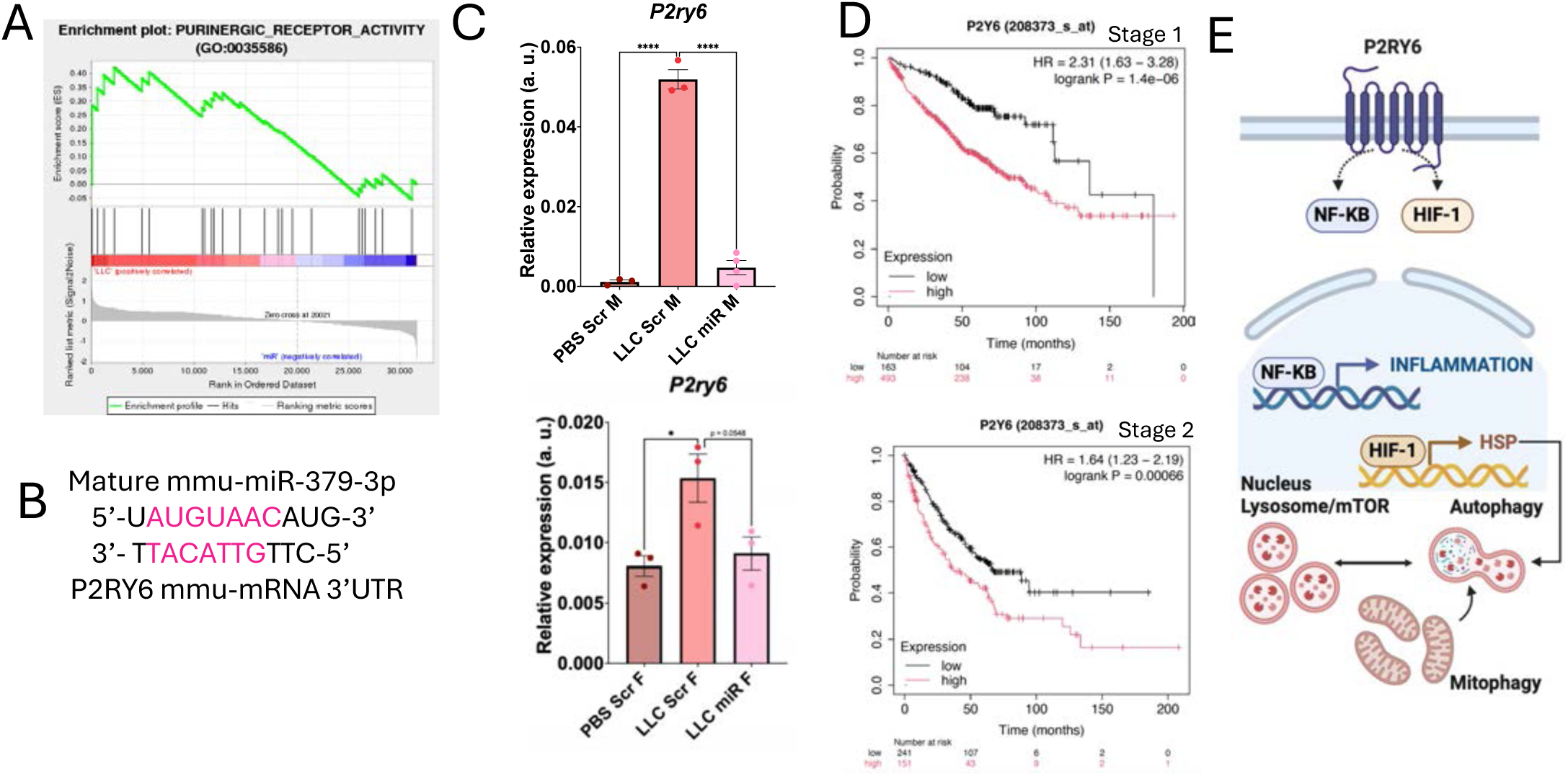
miR-379-3p regulates expression of P2ry6, purinoreceptor associated with inflammation and Ca^2+^ homeostasis. **A.** Gene set enrichment analysis (GSEA) revealed negative association of purinergic receptor activity in LLC miR vs LLC, a predicted target of miR-379-3p. **B.** Murine miR-379-3p seed sequence alignment with 3’UTR of P2ry6. **C.** Regulation of P2ry6 expression in cachectic and cachectic mice treated with miR-379-3p; *P2ry6* relative to *b*2M (*b*2microgloulin) expression was analysed using ΔΔCT. **D.** Survival curves generated from Kaplan-Meier plotter showed that high Pr26y expression was associated with worse prognosis in patients in stage 1 and 2. **E.** Schematic hypothesising P2ry6 link to inflammatory and autophagy signalling.

**Figure 9.**
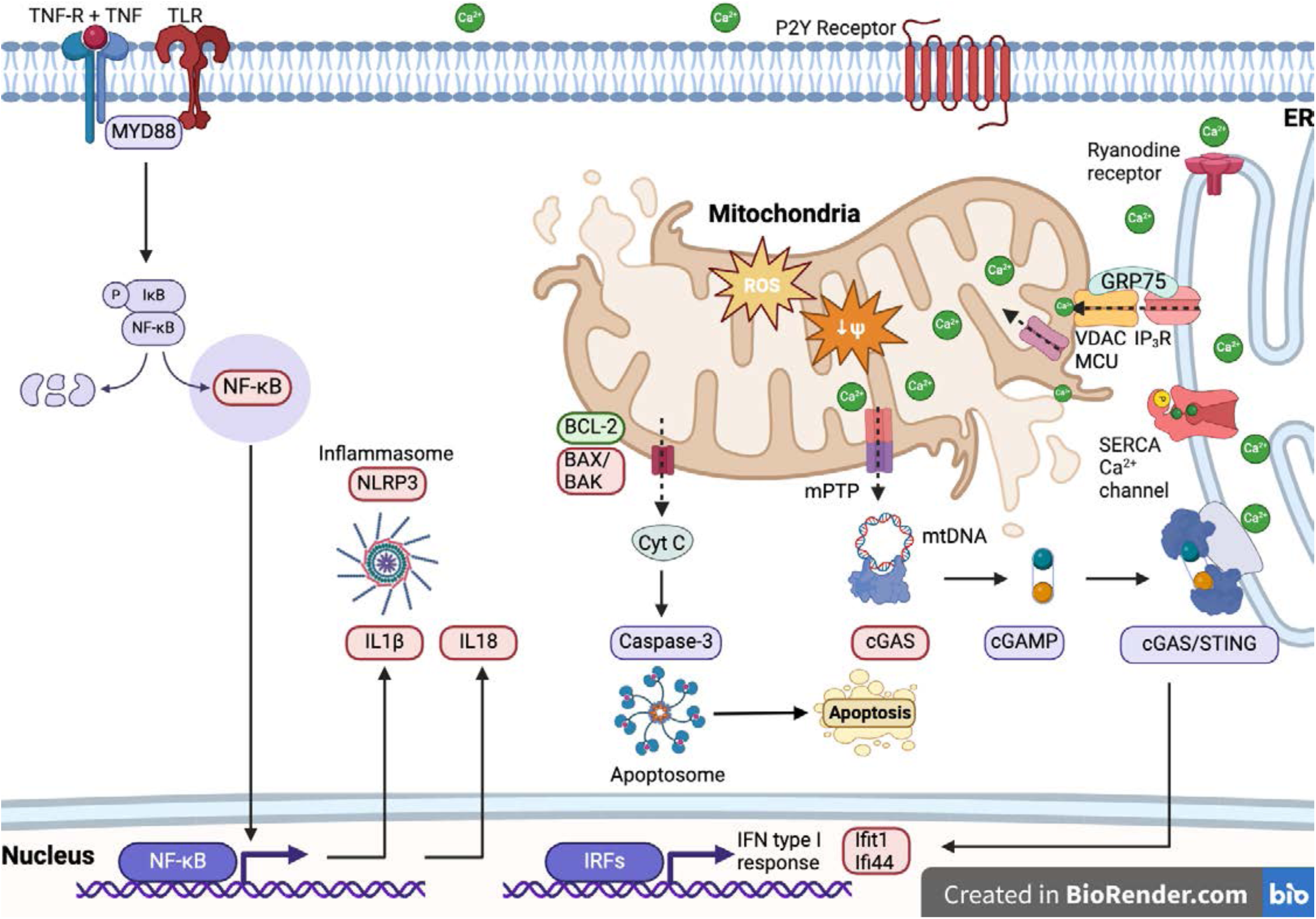
Schematic representation of miR-379-3p as a key regulator of inflammation, mitochondria and apoptosis-related processes in cancer cachexia. Purinergic signalling activates inflammatory pathways; these in turn activate NF-kB further amplifying the inflammatory phenotype. Calcium-regulatory proteins are dysregulated in cachexia, and may contribute to mitochondrial damage and DAMPS, activating cGAS, leading to activation of interferon response, amplifying inflammatory phenotype. Mitochondrial damage and increase in pro-apoptotic factors, likely localising on mitochondrial membrane, may lead to cytochrome release and Caspase activation, and finally apoptosis. miR-379-3p regulates the levels of P2ry6, potentially inhibiting inflammatory cascade. miR-379-3p also regulates the levels of mitochondrial, apoptotic, and inflammatory proteins.

### miR-379 regulated the expression of pyrimidinergic receptor P2ry6

We next investigated potential targets of miR-379-3p. Among target genes conserved between human and mice and regulated in cachexia, was P2ry6 encoding for purinergic receptor P2YY6 and GSEA analysis of female cachectic mice treated with miR-379-3p revealed negative enrichment with purinergic signaling (Fig.8A,B). The aberrant expression of P2RY6 is frequent in human cancers and causes immune evasion [44, 45]. Deletion of *P2RY6* from tumors with high P2RY6 expression inhibits tumor growth, whereas global deletion of *P2ry6* from mice does not compromise viability but improves glucose metabolism and insulin tolerance along with reducing peripheral inflammation [46]. P2RY6 activates the synthesis of prostaglandin E2, a mediator of inflammation and TLR-induced inflammatory responses, and downstream NF-kB activation. We therefore investigated the expression of P2ry6, which was upregulated during cachexia in muscle of male and female mice and downregulated after miR-379-3p treatment (Fig.8C). Survival analyses showed that in lung cancer (LUSC) patients, high levels of P2ry6 were associated with lower survival rates (Fig.8D), supporting P2ry6 as biologically relevant target of miR-379-3p. P2Y6 has been shown to regulate pro-inflammatory signaling, HSF1 activation [44] and calcium-related pathways [47](Fig.8E).

In summary, here we show that early symptomatic cachexia is characterized by changes in mitochondrial proteins, predominantly in male mice, and strong inflammatory signature in female mice. We show that miR-379-3p, levels of which are associated with survival of lung cancer patients, regulated P2ry6 receptor, an upstream regulator of TLR signaling and inflammation. Moreover, we show reduced mitochondrial content and function, potentially associated with mitochondrial damage and cGAS upregulation, linking the mitochondrial phenotype with inflammation in cachexia. Finally, we demonstrate that miR-379-3p coordinately regulates these changes, placing it as a key regulator of cachexia and potential novel therapeutic target.

## Discussion

The precise mechanisms underlying cancer cachexia are not fully understood, however inflammation is a hallmark of CC [1, 2]. It is becoming apparent that changes in muscle, especially at the molecular level, are present before the onset of cachexia phenotype [15]. These include mitochondrial alterations, potentially as adaptive responses to metabolic stress [3, 48]. Patients with no evident symptoms of cachexia present a tendency to experience fatigue during endurance tasks, and an impairment of muscle oxidative metabolism observed in individuals at the pre-cachexia stage of cancer [49]. Therefore, muscle dysfunction can occur before substantial weight loss becomes apparent, highlighting the importance of early intervention strategies aimed at preserving muscle strength and function in cachexia management [50]. These findings underscore the rationale of this study, examining changes in skeletal muscle at the very early symptomatic stages of CC in an LLC mouse model, including differences between males and females. We found limited phenotypic changes at 3 weeks from tumor cell injections (Fig.1), more pronounced in female mice. However molecular changes in muscle from both male and female cachectic mice were observed by global proteomics (Fig.1D, E) and transcriptomics (Fig. 4A-D). Muscle of male mice was predominantly characterized by changes in mitochondrial proteins and energy metabolism, whereas muscle of female mice was characterized by activation of type II interferon response and changes in mitochondrial proteins associated with redox stress, as well as other stress-related pathways, *e.g.*, heat shock proteins.

A set of dysregulated proteins/genes was regulated by miR-379-3p, which is downregulated in cachectic muscle from mice and humans (Fig. 2B, [29, 35]), High levels of miR-379-3p were associated with better survival at early stages of lung cancer (Fig.2A), whereas high levels of its target, P2ry6 (Figs.5,7), regulated in muscle of cachectic mice, were associated with lower survival of patients with lung cancer (Fig. 5D).

miR-379 role during cachexia remains unknown, with few studies reporting miR-379 downregulation in mice and human muscle in CC. Previous studies have focused on the role of miR-379-5p, rather than -3p, that functions in non-small cell lung cancer (NSCLC) [51] and gastric cancer (GC) [52] *in vitro*, focusing on pri-miR-379 overexpression (containing the two strands). miR-379-5p is predicted to target COX-2 and Cyclin B1, and systemic delivery of exosome-encapsulated miR-379-5p showed therapeutic effects *in vivo* in mouse models, highlighting its potential for cancer therapy [53, 54]. In muscle, miR-379-5p, but not -3p, was shown to regulate mitochondrial homeostasis [55].

Here, miR-379 overexpression in mice treated with LLC cells had a protective effect against muscle wasting in the context of cancer cachexia: miR-379-3p preserved muscle mass, fibers atrophy and remodeling and slowed down deterioration of grip strength. Although miR-379-3p showed similar effect on mitochondria and inflammation-related genes and proteins, these effects were observed with variable strength in males and females. A shift from glycolysis to oxidative phosphorylation was observed, indicating enhanced mitochondrial efficiency in males. This was evidenced by the upregulation of mitochondrial proteins such as MT-CO2 and NDUFA2, alongside the downregulation of glycolytic enzymes and type I muscle fibers proteins like MYH7. The results suggest a novel role for miR-379-3p in modulating mitochondrial efficiency and reducing reliance on glycolysis, particularly in males. In females, miR-379-3p reduced proteins involved in the interferon-gamma (IFN-γ) response, which is known to drive pro-inflammatory cytokines such as MyD88 and IL-18 [56], and was validated by qPCR (Fig.6E). This reduction in inflammatory mediators was consistent with decreased activation of pathways like TNF-α and TLR-4 (Fig.6B,E), suggesting that miR-379-3p could alleviate inflammation, a key feature in cachexia. This aligns with earlier findings, where females showed an inflammatory response, particularly in the IFN-γ pathway, while males exhibited a reduction in mitochondrial-related pathways (Fig.1D,E). The effect of miR-379-3p on inflammation and mitochondrial dysfunction suggest a sex-specific impact on cachexia, with females benefiting from reduced pro-inflammatory signaling and males from improved mitochondrial function. While observed limited changes in proteins associated with autophagy or mitochondrial biogenesis, suggesting that these may not be key early events in cachexia and not key targets of miR-379 (Fig.S5) and other models [20].

On the other hand, miR-379-3p may also reduce mtDNA release, triggered by mitochondrial permeability transition pore (mPTP) opening and the pro-apoptotic actions of Bax and Bak, leading to the release of damage-associated molecular patterns (DAMPs) like cytochrome c and mtDNA [57, 58]. miR-379-3p upregulated the anti-apoptotic gene Bcl-2 and downregulated pro-apoptotic genes such as Bax and Bak (Fig.7B) and decreased the abundance of cleaved CASP3, indicative of reduced apoptosis (Fig.7D).

Finally, we associate mitochondrial damage and inflammation, regulated by miR-379-3p with its target gene: P2r6y highlighting a not well understood role of purinergic signalling in cancer cachexia. P2RY6 is frequently dysregulated in cancer and associated with immune evasion [45]. Global deletion of P2Y_6_R improved glucose metabolism and insulin tolerance along with reducing peripheral inflammation [46]. P2Y6 is an activator of prostaglandin E2 and TLR-induced signalling, which initiate inflammatory responses [44]. P2RY6-mediated signalling enhances the levels of TLR-induced pro-inflammatory cytokines and leads to NF-kB activation. The expression of heat shock proteins is strongly induced by inhibition of P2RY6-mediated signalling. Together, this suggests that disrupted levels of P2ry6 in muscle of cachectic mice, may be a key regulator of inflammatory and mitochondrial changes in the early stages of cachexia and miR-379-3p overexpression can ameliorate these changes. We propose that miR-379-3p acts as a regulatory hub for expression of genes associated with mitochondrial content and function, including those regulating mitochondrial integrity and Ca^2+^ signaling between different cellular compartments. miR-379-3p regulates inflammatory response at many levels, possibly through regulation of its target gene, encoding for a purinergic receptor, P2ry6, signaling important in cancer [59]. miR-379-3p prevents muscle loss in cachexia and represents a novel therapeutic avenue for this condition.

### Limitations of the study

Male mice showed limited tumor growth compared to female mice, hence the differences in the effects of cachexia on molecular signature in muscle between males and females may be a result of biological differences between the sexes or the difference in tumor size. Nevertheless, mitochondrial changes likely precede inflammation in early stages of cachexia. Furthermore, miR-379-3p delivery was through intravenous injection, therefore rescue of the muscle phenotype by miR-379-3p, which inhibited tumor growth, particularly considering its target gene: P2r6y expressed in tumor itself, may be a combination of changes within the muscle, as well as the effects of tumor on muscle. Nevertheless, miR-379-3p shows a strong potential as a novel therapeutic candidate to ameliorate cachexia and potentially tumor development. Future studies should validate the role of P2Y6 in cachexia development.

## Methods

### Kaplan-Meier Plotter Database Analysis

The Kaplan Meier plotter was used to assess the correlation between the expression of miR-379 and P2ry6 nd survival in 35k+ samples from 21 tumor types [60]. Gene expression data and overall survival (OS) information were downloaded from GEO, EGA and TCGA databases. The database handled by a PostgreSQL server, integrated gene expression and clinical data simultaneously. To analyze the prognostic value of miR-379/P2ry6, the patient samples were split into two groups according to various quantile expressions of miR-379/P2ry6. The two patient cohorts were compared by a Kaplan-Meier survival plot, and the hazard ratio with 95% confidence intervals and log-rank P value were calculated.

### Study approval and experimental model

All animal studies were approved by the University of Galway Animal Welfare Body and HPRA and maintained as approved under approval AE19125/P091. C57BL6/J young male and female mice (3 months old) were obtained from Charles River, UK. For tumor induction, animals were randomly allocated to groups: PBS Scr n = 3, LLC Scr n = 4 and LLC miR n = 4 for males (M) and females (F), each, and injected subcutaneously, into the left and right flank, with Lewis Lung Carcinoma (LLC) cells (0.5×10^6^ in 100 uL on each side ) or an equal volume of sterile PBS. miR-379-3p overexpression was achieved *in vivo* by intravenous injection (IV) delivery of miR-379-3p mimic or scrambled RNA (Scr) twice: on day 0 and day 7 (D0 and D7), at the dose of 2mg/kg (Table S1) [20].

### Tissue collection and grip strength

Mice were sacrificed by anesthetic overdose 3 weeks after the first injection. Tissues were immediately collected and frozen at -80 for sample processing (RNA and protein isolation) or placed in OCT and snap-frozen in isopentane for histological analyses. Body weight (BW) and grip strength (GS) were measured at days 1, 14 and 21. Tumor and muscle weights were measured at D21 at the end of the experiment. GS measurements were taken using a grip strength meter (GSM-Ugo Basile 47200). Each GS measurement session was performed at the same time, and by the same operator. Briefly, the mouse was then allowed to grasp at the cross bar with the forelimbs and pulled backwards gently by the tail, until the release of the bar. The data collected was the corresponding force and time. A total of three attempts were given. The maximum force was recorded for each experiment and normalized to BW. Change over time was calculated as follows: normalized force final (day 21) - normalized force initial (day 0).

### Histology

Ten μm sections of GAS was stained with Fluorescein-labelled wheat germ agglutinin, WGA, staining, to visualize myofibers size as before [20]TA muscle was used for Hematoxylin and Eosin (H&E) staining as before [37]. Sections were mounted using Hydromount mounting medium. Minimal Feret’s diameter was measured using ImageJ, as described previously [61]. TA was used for myofiber type staining [61]. Sections were washed in PBS and blocked in diluted HS and Mouse-on-Mouse (MOM) blocking agent. Next, sections were incubated with anti-MHC I, - MHC IIa and -IIb 1:250 in diluted HS. Secondary antibodies: Alexa fluor 350, 488 and 532 were diluted 1:500. Images were taken using EVOS Imaging system M7000. Analysis was performed with Image J software.

### RNA and protein isolation

PreOmics BeatBox® was used for tissue homogenization of 10 mg of QUAD for 10 minutes. Protein was isolated after homogenization following iST 96x (PreOmics) 96-sample kit following manufacturer indications until lyophilization and samples were analyzed by the proteomics facility NICB at Dublin City University. Muscle homogenate was processed using miRvana isolation kit following the manufacturer’s protocol. NanoDrop2000 was used for determination of sample purity and concentration.

### Global mass spectrometry

Quality controls were performed for proteome analysis. The quality control assessment included density plots, correlation heatmaps, boxplots, and PCA to ensure consistency in expression data across samples. R software was used to generate principal component analysis plots (PCA) and quality control plots (density, boxplot and heatmap).

Mass spectrometry raw files were processed using MaxQuant (v 2.4.10.0) against a *Mus musculus* protein database (UniProt, proteome ID: UP000000589, taxonomy ID: 10090, downloaded May 2024). Fixed modifications included carbamidomethylation of cysteine, while variable modifications included oxidation of methionine and acetylation of protein N-termini. Trypsin was specified as the proteolytic enzyme, allowing up to two missed cleavages.

The minimal peptide length was set to 7 amino acids. Peptide and protein identifications were filtered at a 0.01 false discovery rate (FDR) at the PSM, protein, and site levels. Label-free quantification (LFQ) was enabled. Results were filtered at 0.01 false discovery rate (FDR) at both peptide and protein levels. The “proteinGroups.txt” file obtained from MaxQuant was analyzed using Prostar software [62].

### RNA sequencing

For p-value, represented as -log10(p value), cut off was selected in 0.05 and log 2 fold change threshold of 1.3 as previously [63], representing approximately a 2.46 times change in expression. R script software was used (R 4.3.2 and the package ggplot2 3.4.4) [64, 65] for creating volcano plots. KEGG and GO Terms enrichment analysis and dot-plots were generated with R studio ggplot2 library (Ge et al., 2020). Gene interaction networks were created in STRING Software and K-means clusters. Pathway analysis of RNA-seq data was performed using PathVisio 3.3.0 [66]. The GSEA analysis was performed to identify gene sets significantly enriched in two phenotypes, using RNA-seq data. The dataset contains 31,649 features (genes). Size filters were applied (min=15, max=5000). Scripts: https://github.com/mariaborjaag/Scripts

### RT-qPCR

miR cDNA synthesis using 10ng RNA was performed using miRCURY LNA RT kit and qPCR was performed using miRCury qPCR kit according to manufacturer’s protocol as before [67]. miR relative expression was normalized to Snord68 and Unisp6 RNA and calculated using delta-delta Ct method for *in vivo/in vitro* miR expression assessment. For gene expression in male and female mice, cDNA synthesis (mRNA) was performed using 500 ng RNA and SuperScript II. qPCR was performed using SybrGreen qPCR as described before [27]; Expression relative to *S29* housekeeper gene was calculated using the delta-delta Ct method. Primers are listed in Table SX.

### miR-379-3p target prediction

mmu-miR-379-3p seed sequence was obtained from miRbase (https://www.mirbase.org/). Significantly downregulated genes and proteins were input into ENSEMBLE Bio Mart software to obtain 3’UTR transcript sequences which were compared to miR-379-3p seed sequence. Venn diagrams were created in R studio software. Predicted target screening was performed using TargetScanMouse and Human 8.0 software [68].

### Western Blotting

Protein lysis and Western Blotting were performed as described previously [20]. Briefly, homogenized protein lysates were diluted in Laemmli buffer, and 30 µg of protein were loaded and separated on 6-14% SDS PAGE gels. Proteins were transferred to nitrocellulose membranes using a semi-dry blotter and stained with Ponceau-S. Membranes were blocked in 5% milk or 5% BSA in TBS-T for 1 h at room temperature. Next, membranes were washed in TBS-T and incubated with primary antibodies (listed in Table SX) at a dilution of 1 in 1000 in blocking buffer, antibody details listed in Table 642. Li-Cor bioscience anti-rabbit/mouse secondary antibodies were diluted 1 in 20,000 in TBS-T and visualized using Li-Cor Biosciences Odyssey Fc. Western blot images were quantified using Image J (version 1.51, NIH, USA: https://imagej.nih.gov/ij/).

### Cell culture

C2C12 myoblasts and LLC cells were cultured as before in GM [69]. Myogenic differentiation was induced by placing 90% confluent cells in DMEM supplemented with 2% horse serum (HS) and 1% PS (differentiation media; DM). Myoblasts or myotubes were co-culture with LLC cells using transwell system membranes of 0.4 µm which impedes cellular migration. Co-culture was performed in a 2:1 ratio proportion, optimization was performed.

Seeding of C2C12 on cell culture and LLCs on inserts was performed on the same day [70, 71]. Myoblasts/myotubes co-cultured with DM filled inserts were used as control (Ctrl). C2C12s cells were plated at either 80% confluency (MF20 immunostaining) or 50% confluency (Cell probe staining). miR-379-3p/Scr treatment (100nM) was performed at the same time points as transwell co-culture of DM /LLCs for 72 hours.

Myoblast differentiation (myogenesis) and myotube atrophy were examined 5/3 days after co-culture, respectively, by immunostaining with myosin heavy chain: MF20 antibody concentrate (DSHB) as before [69]. Images were obtained using M7000 Imaging Systems. Each group contained 3 replicates, and a total of 5 images were taken per well. Images were analyzed using ImageJ [69].

### Mitochondrial assays

Mitochondrial membrane potential was measured using TMRN™. Mitochondrial content was assessed using MitoTracker™ Green FM. Mitochondrial ROS production was assessed by MitoSOX™. Mitotracker Green, MitoSOX or TMRN probes were incubated for 30 min at 37°C [72]. Mitochondrial membrane potential and mitochondrial ROS production were quantified by the mean fluorescence intensity area of each cell normalized by the mitochondrial content. Mitochondrial content was quantified by the mean of the green channel grey value intensity of each individual cell. Images were taken at 40x/60x magnification using EVOS M7000 Imaging Systems. The fluorescence intensity was quantified using the CTCF formula [73].

Seahorse XF HS Mini bioanalyzer was used to measure the oxygen consumption rate (OCR) in C2C12 cells transfected with Scr or miR-379-3p. C2C12 myoblasts were seeded at 8000 cells each well in an 8-well XF HS Mini Analyzer plate, incubated for 24 hours, and then treated with miR-379. After another 24 hours, the cells were prepared for the Mito Stress Test Assay using specific reagents, including oligomycin, FCCP, and a combination of rotenone and antimycin A. The assay involved calibrating the cartridge, adding the reagents to designated ports, and running the test to determine OCR, which was then analyzed using Seahorse Bioanalyzer software [74]. Oligomycin (2 µM), an ATP synthase inhibitor, was used to determine the proton leak; carbonyl cyanide p-trifluoromethoxy phenylhydrazone or FCCP (2 µM), a mitochondrial uncoupler, will give an indication of the maximal uncoupled respiration and antimycin A (1 µM), a complex III inhibitor, will provide information about the nonmitochondrial respiration. DAPI nuclei staining was used to determine the number of cells.

### Schematics

All schematics were generated in Biorender.

### Statistics and reproducibility

Data are expressed as mean ± standard error of the mean (SEM) or mean ± standard deviation (SD), as indicated in the figure legends. Statistical significance for normally distributed data was determined using Student’s t-tests for comparisons of 2 groups or analysis of variance (ANOVA) followed by Tukey’s test for comparisons of 3 or more groups. Significance was set at *P* < 0.05 unless otherwise stated. Chi-square test was used to compare the distribution into multiple categories.

Statistical analyses were performed with Prism 7 (GraphPad Software), R (v.4.0.5). Quantification of Western blots was performed using ImageJ 1.53a. Kaplan Meier survival plots were made using https://kmplot.com/and analyzed using Log-rank Mantel-Cox test.

## Acknowledgements

Proteomics facility, DCU and Dr. Jose Casas Martinez, University of Galway, Ireland. This work was supported by funding to KGW: Research Ireland SFI19/FFP/6709; IRCLA/2017/101. MBG was supported by funding from Research Ireland: IRC GOIPG/2022/646 and CS is funded by the Chinese Scholarship Council.

## Author contributions

Conceptualization: KGW, BMcD, MBG, RD

Investigation: MBG, RGO, CS, TS, EMcC, CSN

Funding acquisition: KGW, MBG, CS

Writing, review, editing: All authors

---

**Figure S1.**
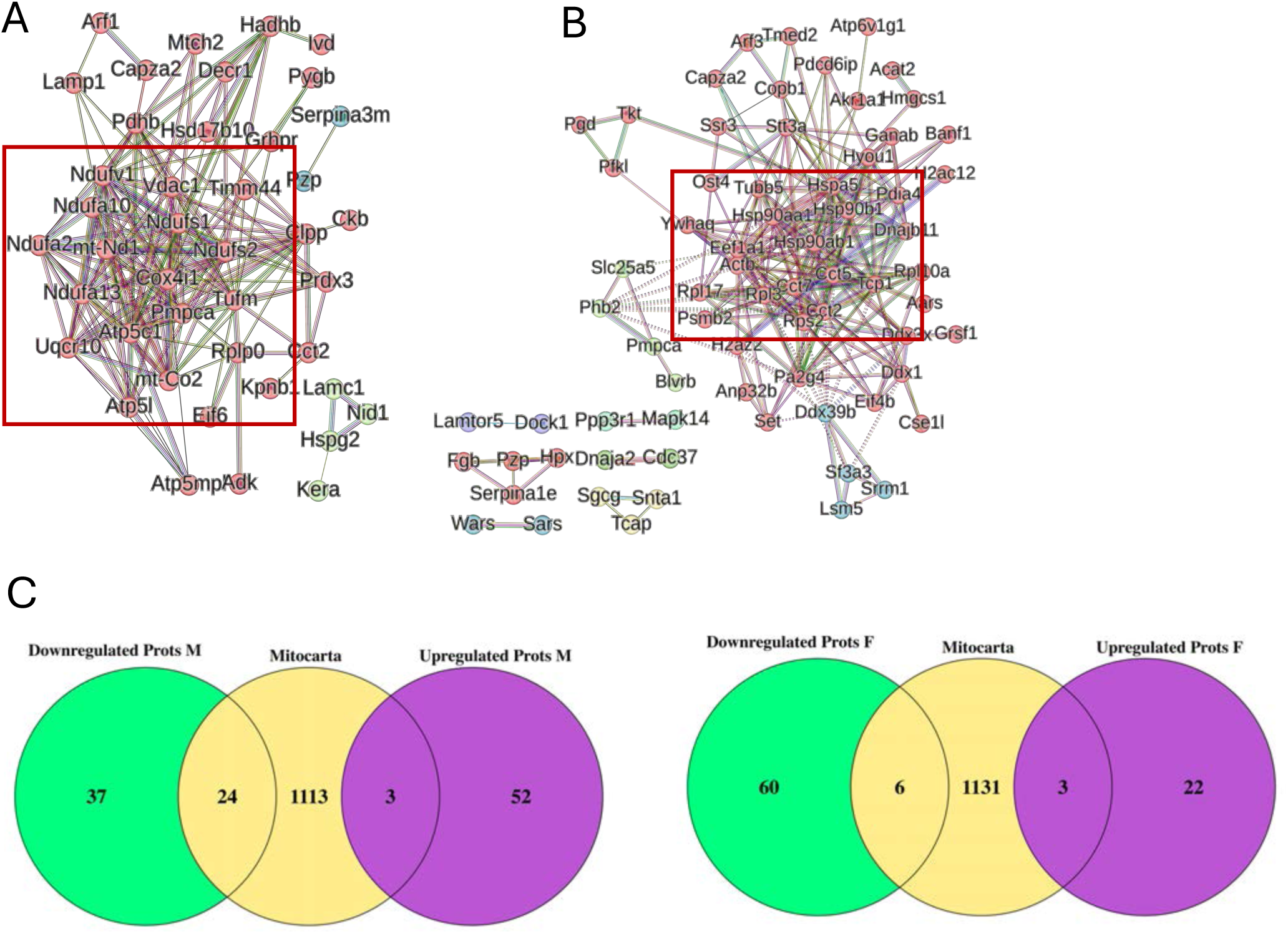
Proteome changes in male and female muscle during early stages of cancer cachexia: mitochondrial dysfunction in males and inflammation and impaired protein folding in females. **A.** Interaction network of downregulated in males was represented, finding 3 clusters related to inflammation (SERPINA3M), extracellular matrix (ECM) (KERA) and ATP biosynthesis, oxidative phosphorylation and cellular respiration (MT-ND1, -CO2, NDUFA2/10). **B.** Interaction network of downregulated in females was represented, finding a highly connected network divided into 3 clusters, mainly associated with protein folding (HSP90AA1, HSP90AB1, ACTB, CCT5, TCP1, and HSPA5), splicing (LSM5, SRRM1, SF3A3 and DDX39B) and mitochondrial protein processing (SLC25A5). Interaction network of more abundant proteins in females was represented, finding 6 clusters, mainly associated with inflammation and haemostasis (PZP, SERPIN3E, MAPK14, FGB, HPX) and stress response (CDC37, DNAJA2). **C.** Venn diagrams show overlap between proteins regulated in muscle of males and females during cachexia and overlapping with MitoCarta3.0 set of genes associated with mitochondria. PBS – control; LLC – cachectic mice; LLC miR-cachectic mice treated with miR-379-3p.

**Figure S2.**
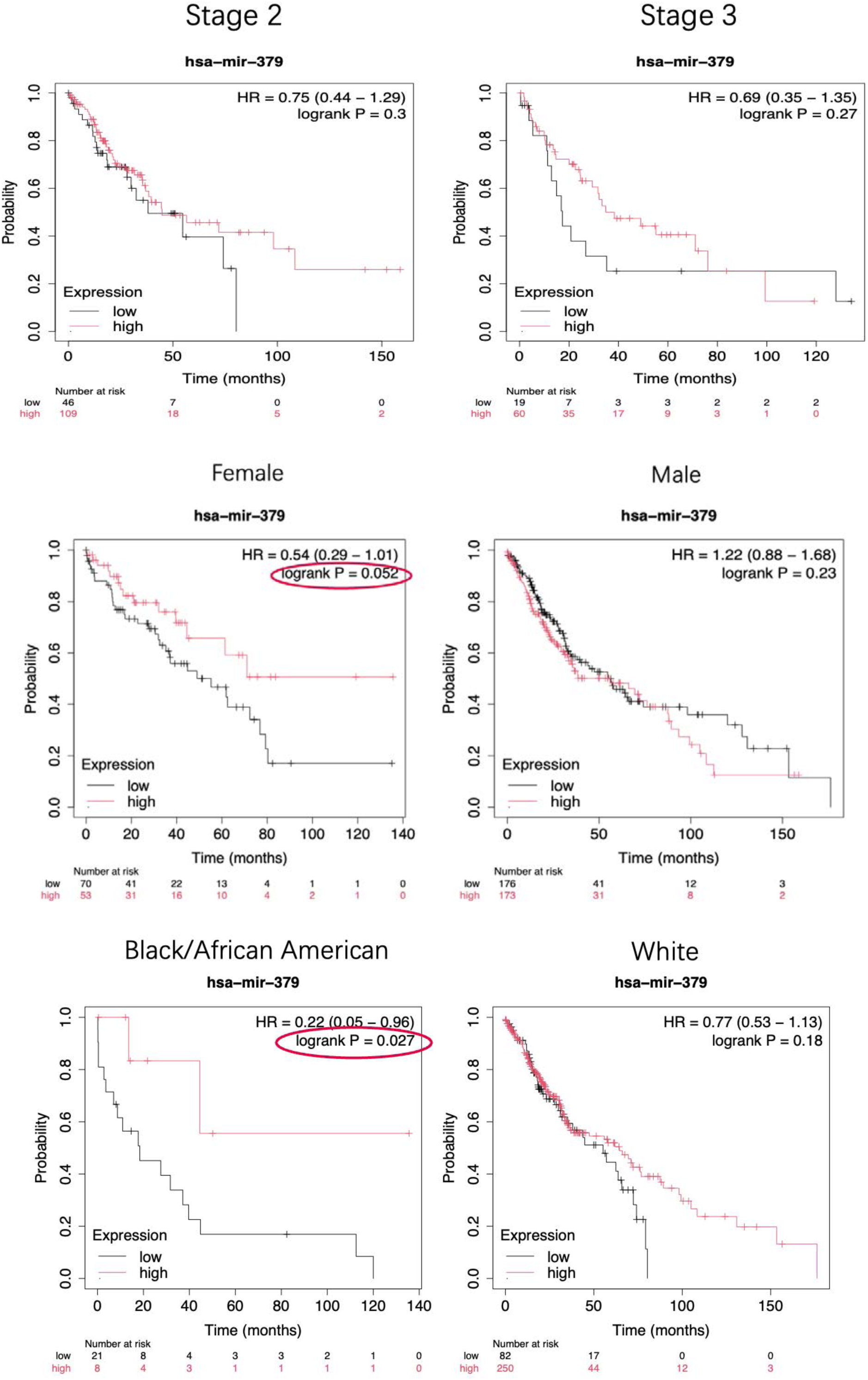
The levels of miR-379 are associated with survival in lung cancer patients. Survival curves generated from Kaplan-Meier plotter showed that low miR-379 expression was associated with worse prognosis in patients at the early stage of lung cancer (HR=0.47, 95% CI= 0.27 to 0.84, P= 0.0084).

**Figure S3.**
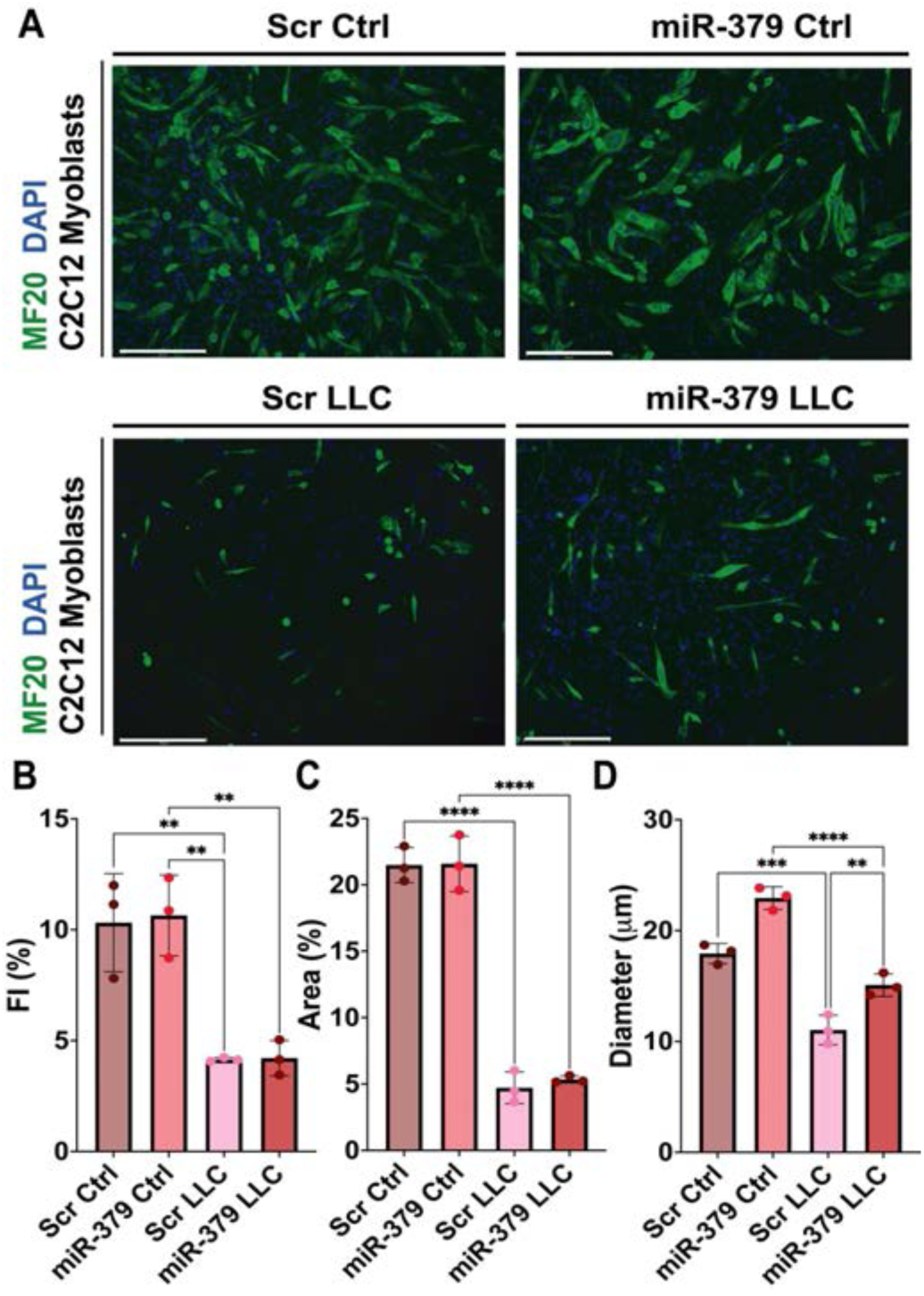
miR-379 did not change myogenic potential in a mouse *in vitro* model of cachexia. **A.** MF20 (anti-myosin heavy chain; green) and DAPI (blue) immunostaining were performed for atrophy assessment. **B.** Fusion index, FI, was measured, and **C.** Myotube area was assessed. **D.** The diameter of myotubes was also analysed. LLC co-culture decreased myotube size, and FI, miR-379 ameliorated this reduction a model of cachexia *in vitro*. Representative images were shown; Error bars showed SD; * p < 0.05; ** p<0.01, *** p<0.001, **** p<0.0001; One-way ANOVA followed by Tukey’s multiple comparison test was performed to assess significances. Scale bar 275 μm. Analysis performed in ImageJ software. miR-379 prevented myotube size loss during cachexia.

**Figure S4.**
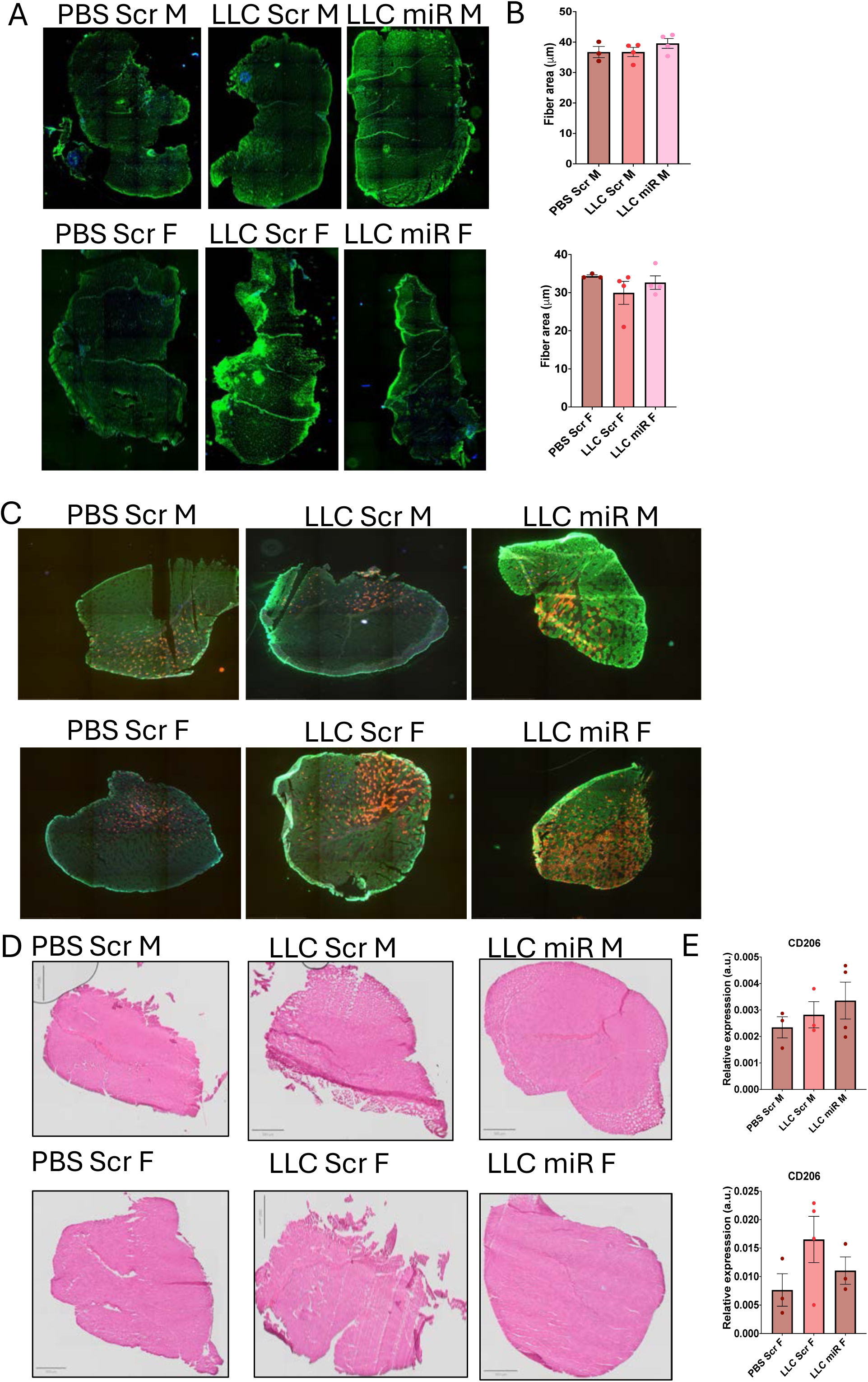
Myofibre size, type and immune cell infiltration in miR-379-3p treated cachectic male and female mice. **A.** Full muscle images of WGA. **B.** Average fibre size was represented per group in males and females. **C.** Full muscle images of fiber type immunostaining. **C.** Full images of H&E staining. **D.** Cd206 relative expression was analysed using ΔΔCT methods and did not show statistically significant differences. Graphs showed relative expression to b2M (b2microgloulin) housekeeper. Error bars showed SEM; PBS Scr, n=3 males and females; LLC Scr, n=4 males and females and LLC miRs, n=4 for males and females. PBS – control; LLC – cachectic mice; LLC miR-cachectic mice treated with miR-379-3p.

**Figure S5.**
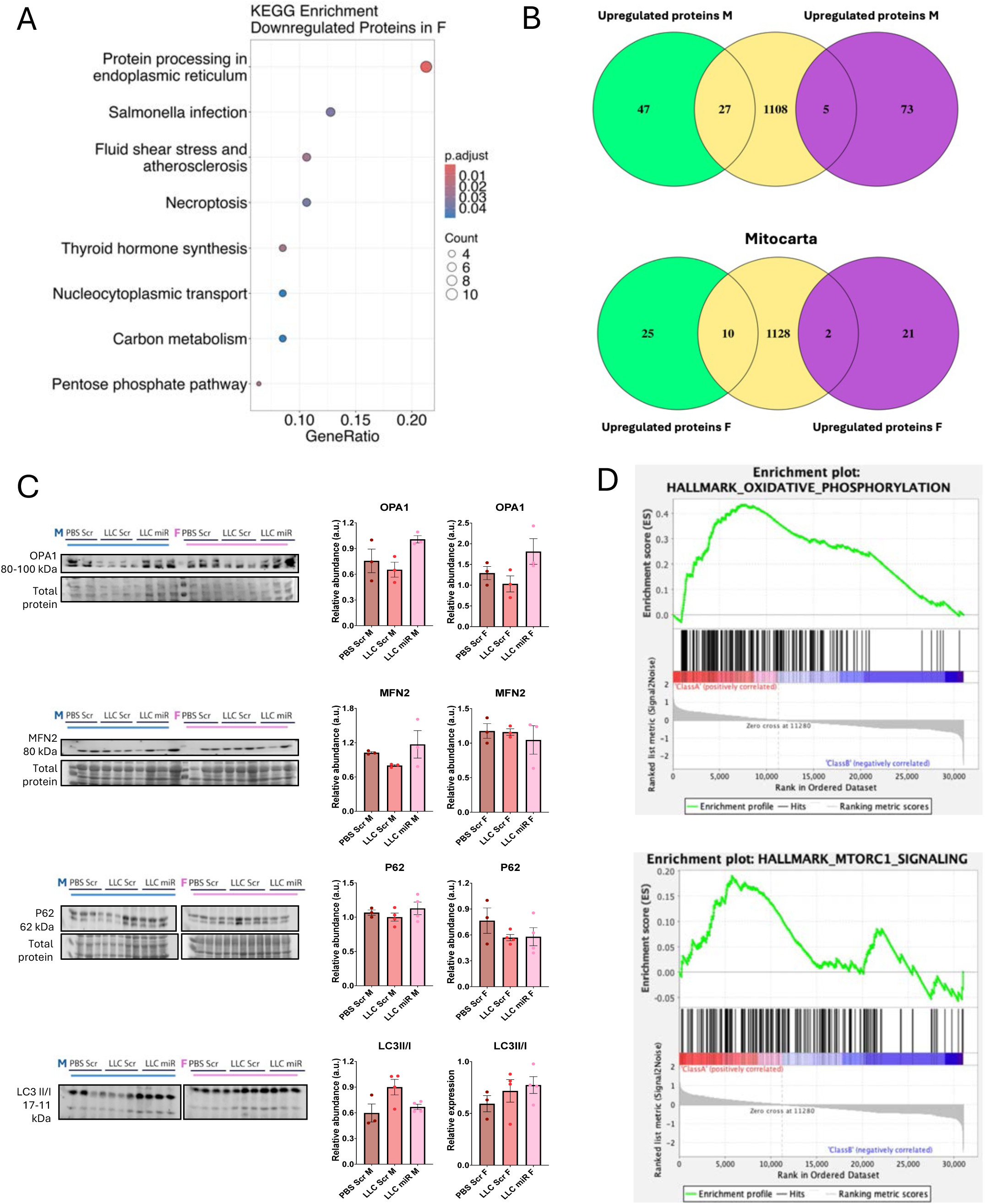
miR-379-3p regulates mitochondrial proteins in cachexia. **A.** KEGG enrichment of proteins downregulated in females. **B.** Venn diagrams were performed in R studio 4.3.2, showing shared proteins (up and downregulated) in the muscle of LLC miR mice compared to LLC mice, with MitoCarta3.0 genes. Upregulated proteins were linked to mitochondrial function and metabolism in LLC miR males and to mitochondrial function and homeostasis in LLC miR females **C.** Western blot was performed to assess protein abundance of OPA1, MFN2, p62 and LC3B, associated with mitochondrial biogenesis, fusion, and autophagy. Their levels were not significantly altered in the muscle of cachectic or cachectic mice treated with miR-379-3p. **D.** GSEA show positive enrichment of mTOR and oxidative phosphorylation-associated genes in female cachectic mice treated with miR-379-3p. Bar graphs show densitometry analysis, performed in ImageJ software. N=3 in each group. In P62 and LC3B WB: PBS Scr (M and F) n=3; LLC and LLC miR (M and F) n = 4. PBS – control; LLC – cachectic mice; LLC miR-cachectic mice treated with miR-379-3p.

**Figure S6.**
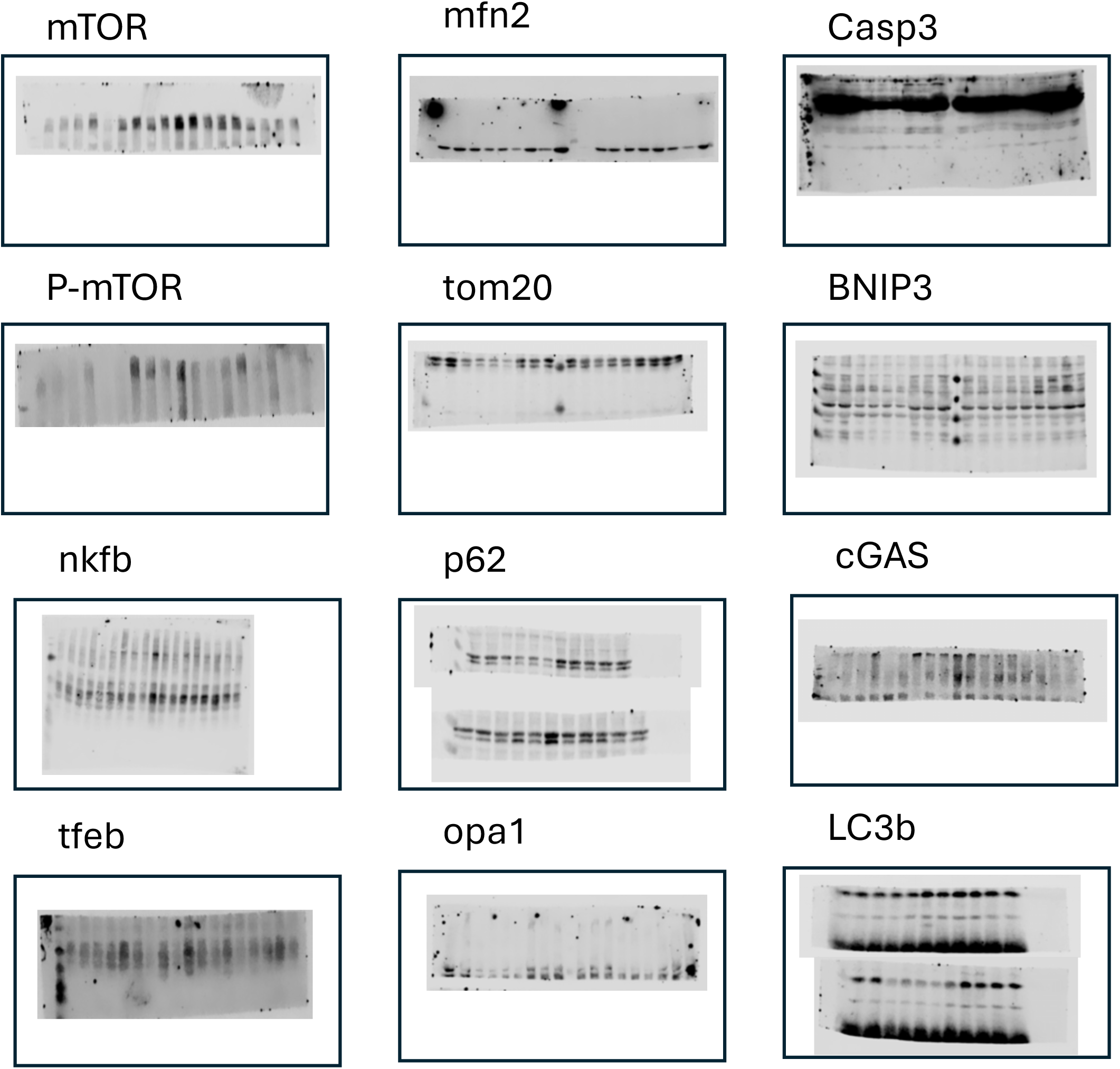
Original WB membranes from Studio Lite software.

**Supplementary figure 7.**
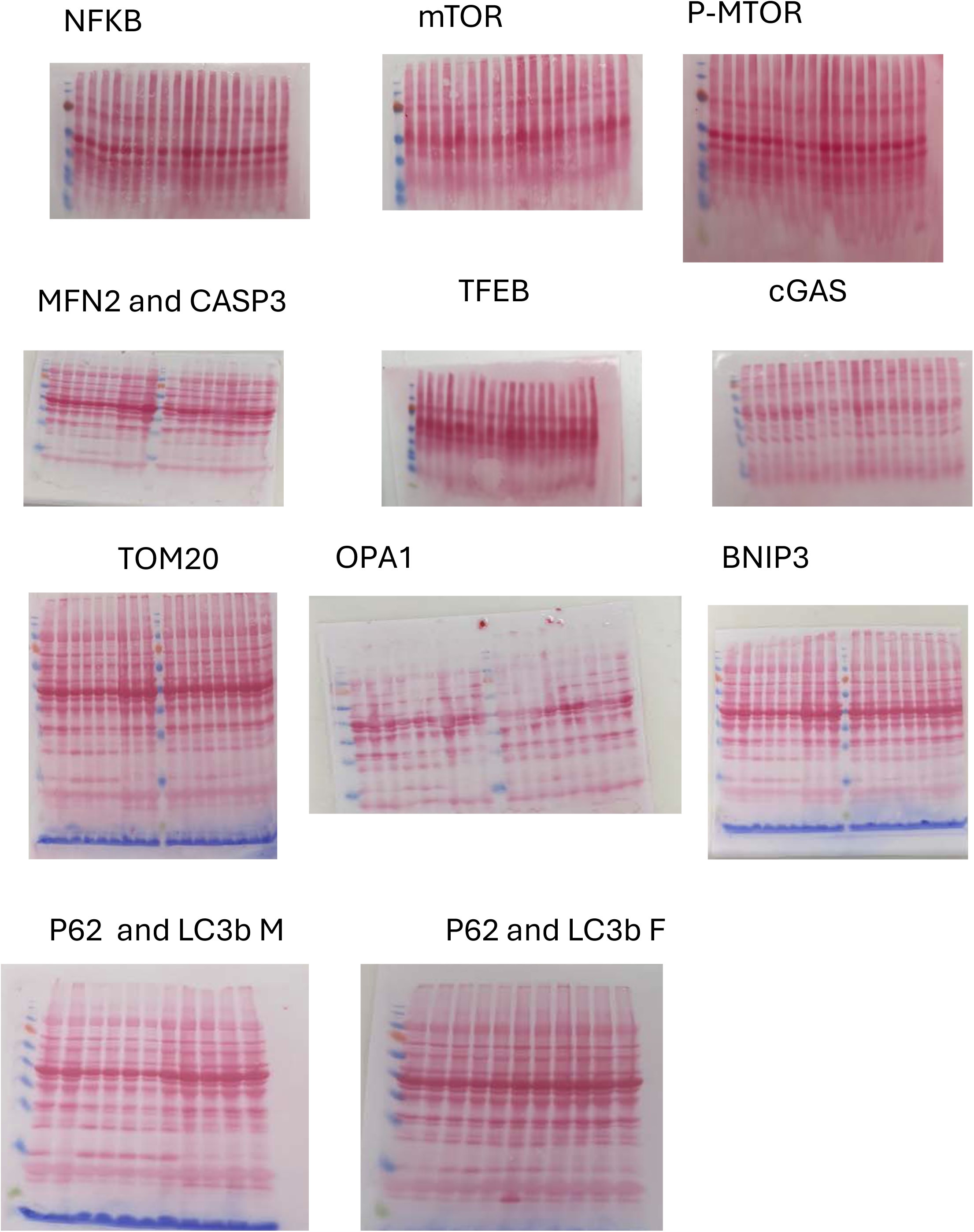

**Figure S8.**
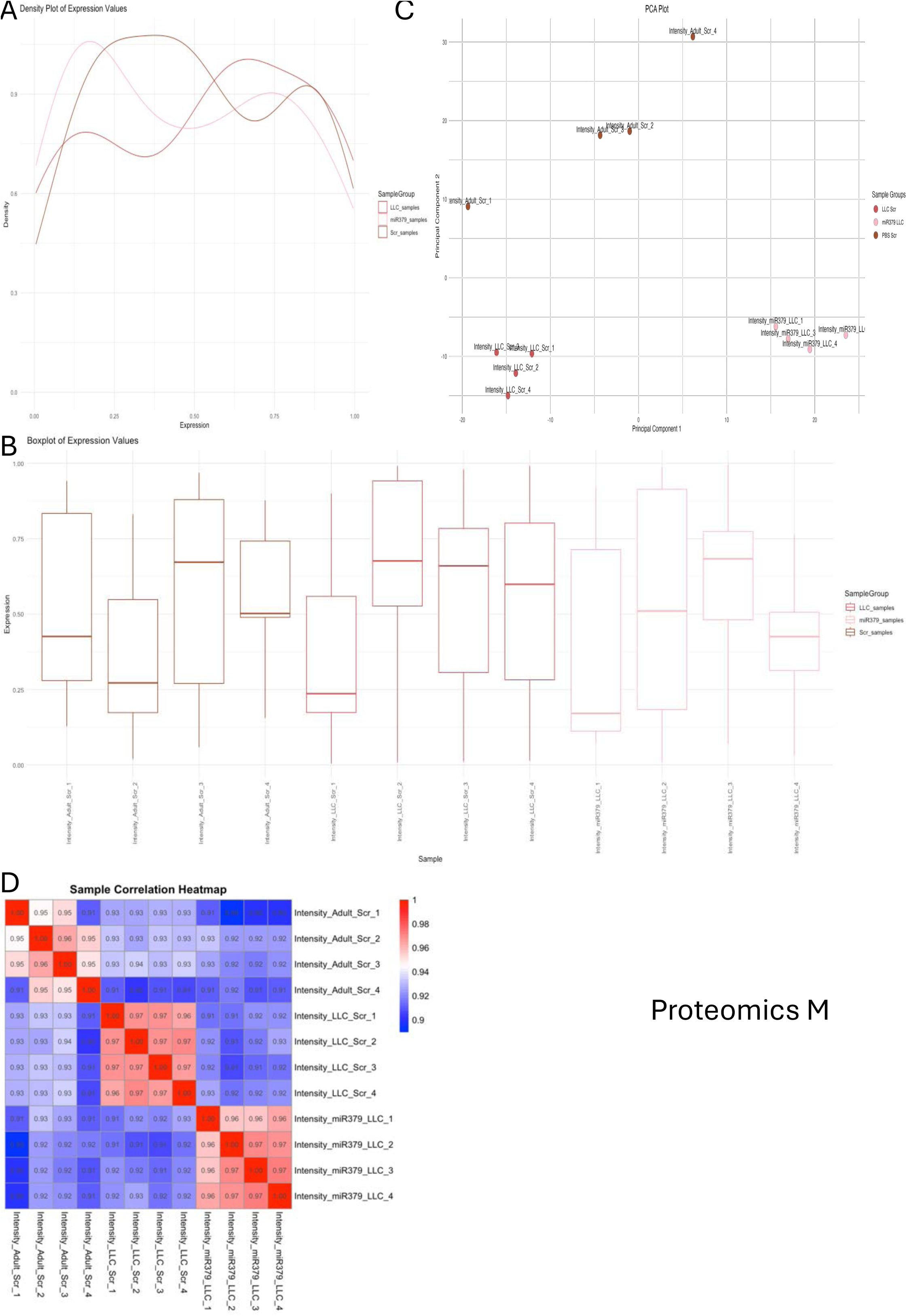
Quality control plots showed proteomics data was of good quality for further analysis in males. **A.** The density plots for PBS Scr (brown); LLC Scr (red) and LLC miR (pink) samples showed some overlap but also distinct peaks, suggesting some differences in expression distributions between the two conditions. **B.** The boxplot provided a summary of the expression values for each sample: the median expression values (lines inside the boxes) and the spread (interquartile range) were shown for each sample. **C.** The PCA showed three clusters, indicating biological separation. **D.** Correlation heat map showed correlation coefficients between the samples. Correlations close to 1 indicate strong similarity. Heat map showed high correlation within groups and low between groups. N=4.

**Figure S9.**
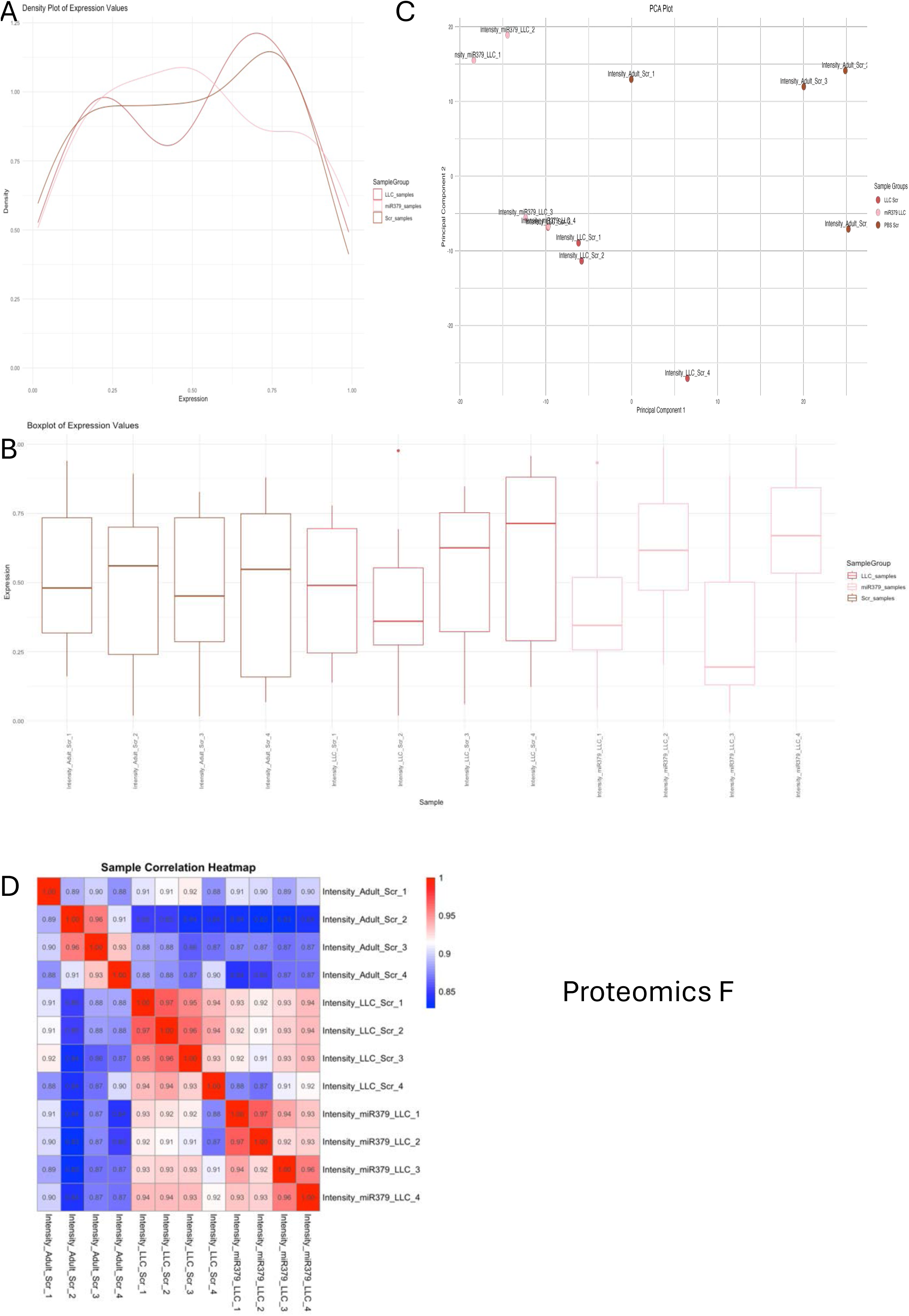
Quality control plots showed proteomics data was of good quality for further analysis in females. **A.** The density plots for PBS Scr (brown); LLC Scr (red) and LLC miR (pink) samples showed some overlap suggesting some similarities in expression distributions between the two conditions. **B.** The boxplot provided a summary of the expression values for each sample: the median expression values (lines inside the boxes) and the spread (interquartile range) were shown for each sample. **C.** The PCA showed distinct clusters. **D.** Correlation heat map showed correlation coefficients between the samples. Correlations close to 1 indicate strong similarity. Heat map showed high correlation within groups and low between groups with some more similarity in LLC and LLC miR samples. N=4.

**Figure S10.**
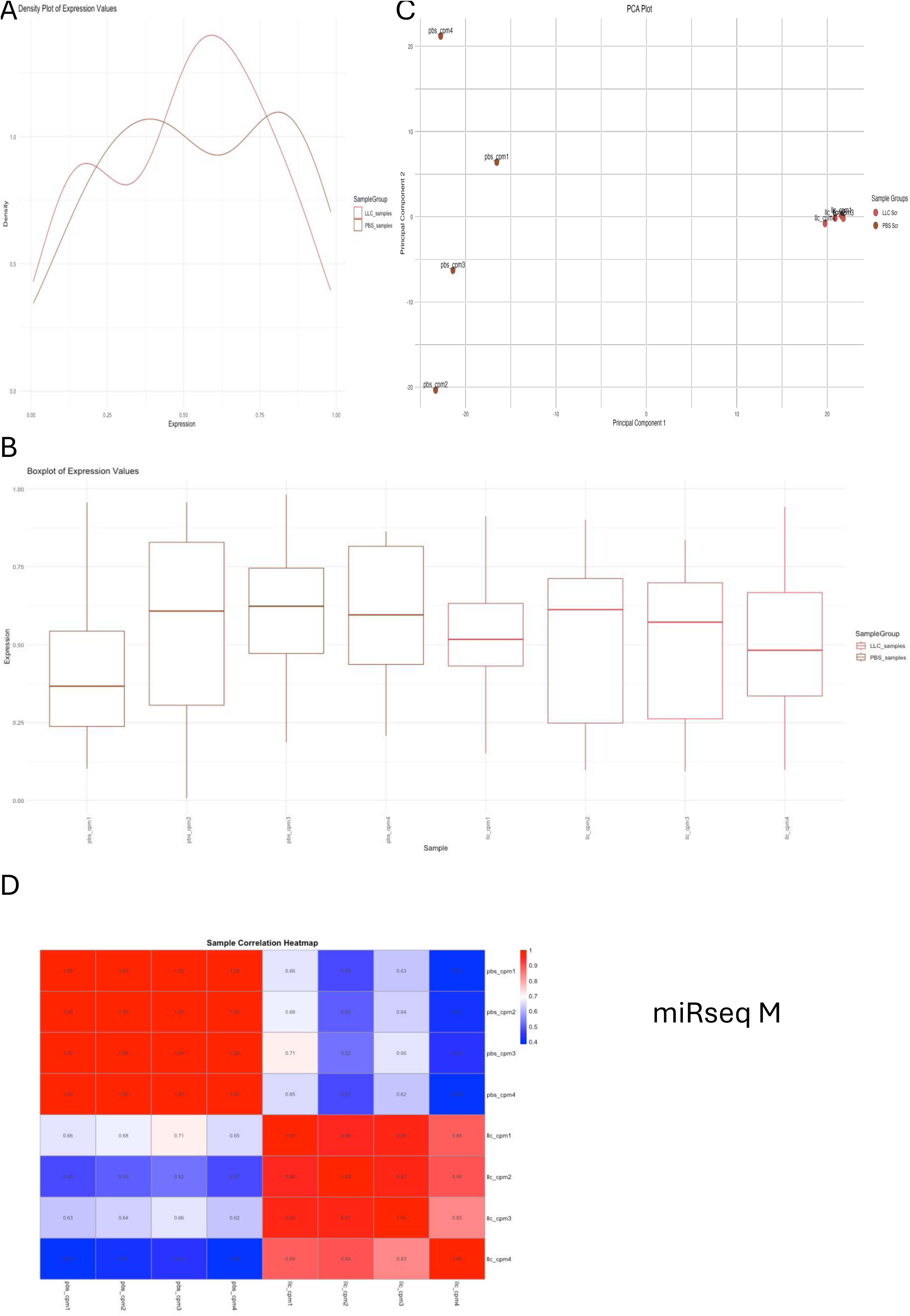
Quality control plots showed miR-seq data was of good quality for further analysis in males. **A.** The density plots for PBS Scr (brown); LLC Scr (red) samples showed some overlap suggesting some differences in expression distributions between the two conditions. **B.** The boxplot provided a summary of the expression values for each sample: the median expression values (lines inside the boxes) and the spread (interquartile range) were shown for each sample. **C.** The PCA showed distinct clusters for LLC and PBS samples. **D.** Correlation heat map showed correlation coefficients between the samples. Correlations close to 1 indicate strong similarity. Heat map showed high correlation within groups and low between groups. N=4.

**Figure S11.**
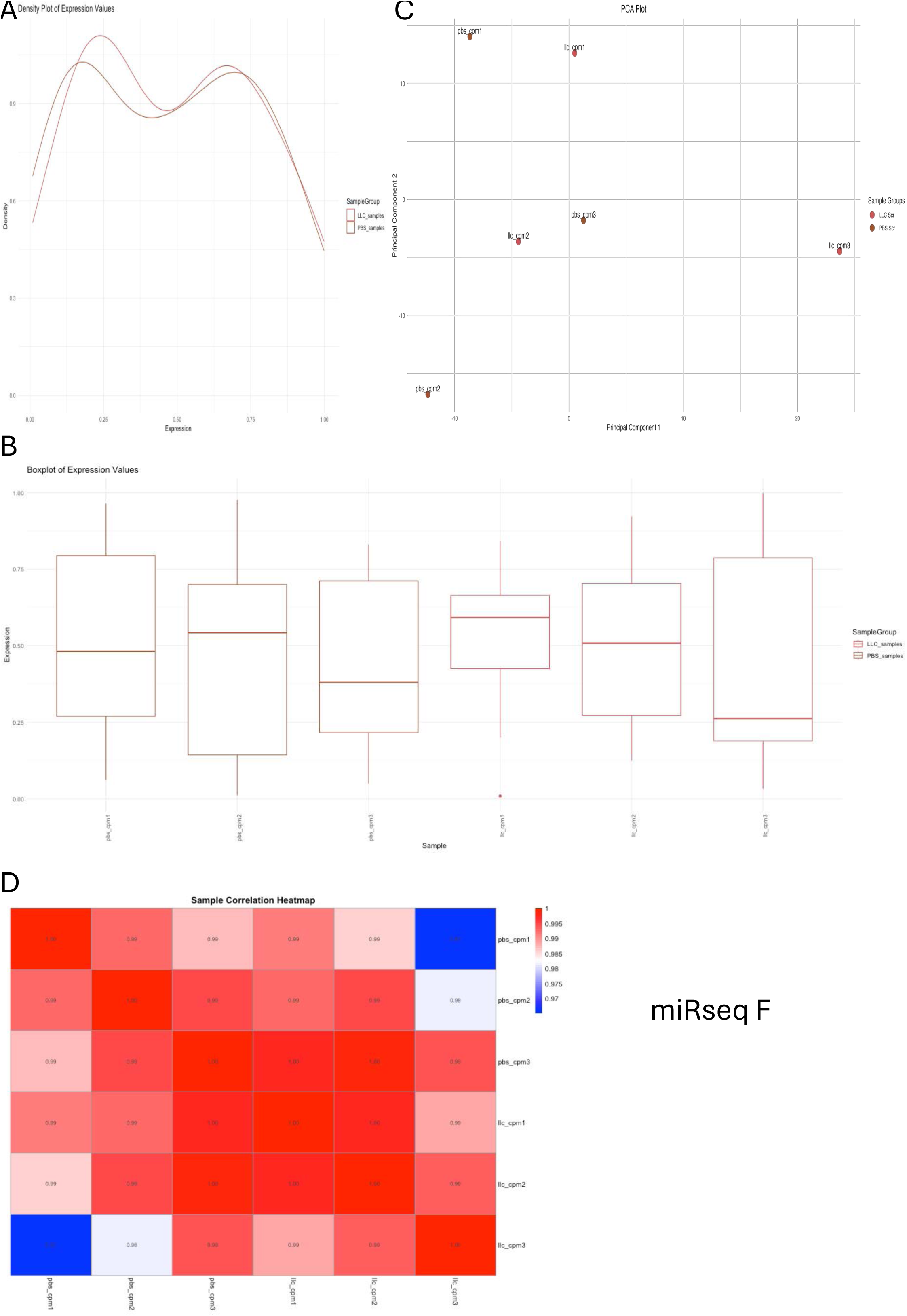
Quality control plots showed miR-seq data some correlation between samples in females. **A.** The density plots for PBS Scr (brown); LLC Scr (red) samples showed some overlap suggesting some similarities in expression distributions between the two conditions. **B.** The boxplot provided a summary of the expression values for each sample: the median expression values (lines inside the boxes) and the spread (interquartile range) were shown for each sample. **C.** The PCA LLC and PBS samples did not cluster. **D.** Correlation heat map showed correlation coefficients between the samples. Correlations close to 1 indicate strong similarity. Heat map showed high correlation among groups. N=3.

**Figure S12.**
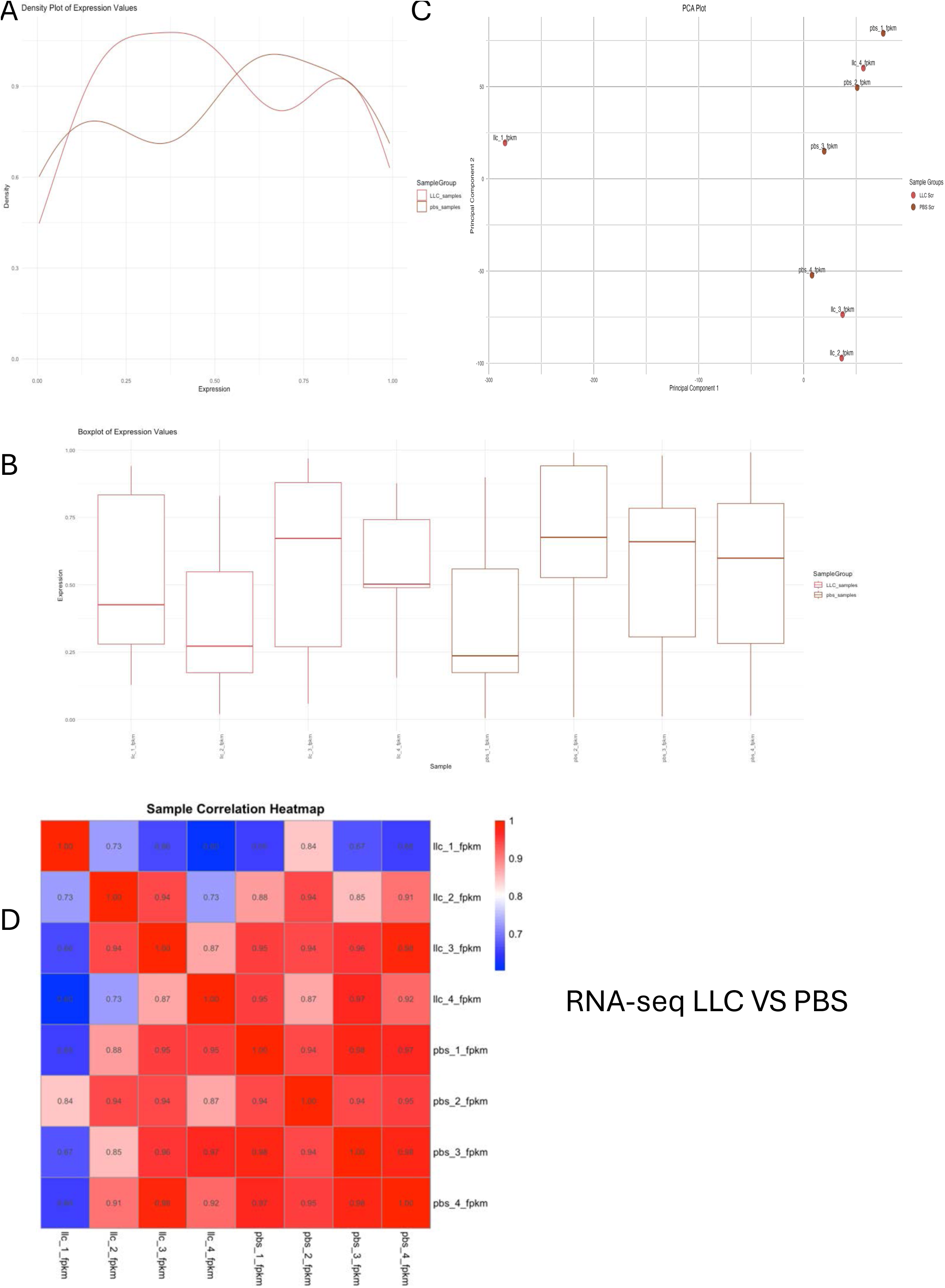
Quality control plots showed RNA-seq data was of good quality for further analysis in females. **A.** The density plots for PBS Scr (brown); LLC Scr (red) samples showed some overlap suggesting some differences in expression distributions between the two conditions and two distinct peaks. **B.** The boxplot provided a summary of the expression values for each sample: the median expression values (lines inside the boxes) and the spread (interquartile range) were shown for each sample. **C.** The PCA showed distinct clusters for LLC and PBS samples. **D.** Correlation heat map showed correlation coefficients between the samples. Correlations close to 1 indicate strong similarity. Heat map showed some correlation between groups. N=4.

**Figure S13.**
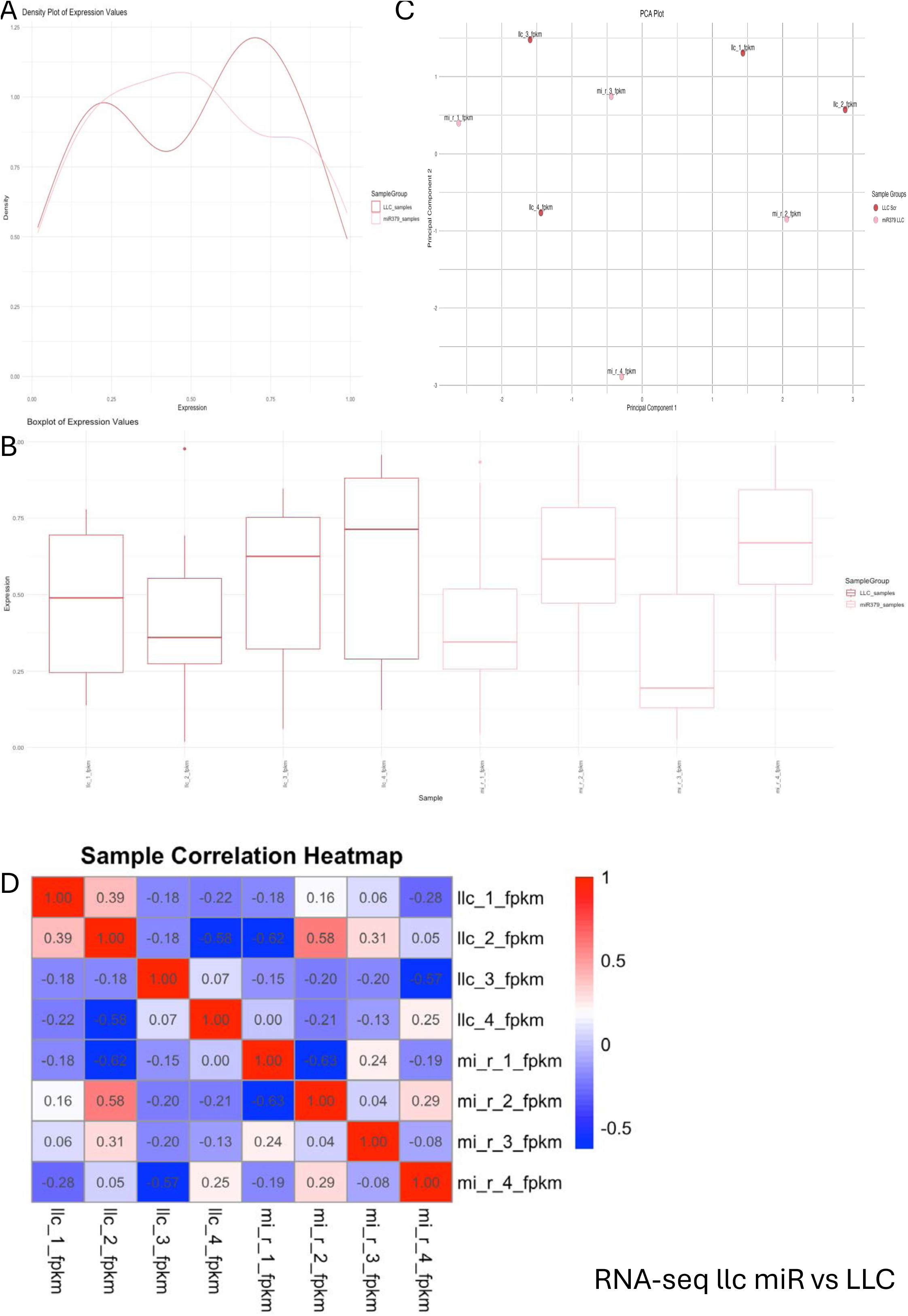
Quality control plots of RNA-seq data in LLC miR females. **A.** The density plots for LLC Scr (red); LLC miR (pink) samples showed some overlap suggesting some differences in expression distributions between the two conditions and two distinct peaks. **B.** The boxplot provided a summary of the expression values for each sample: the median expression values (lines inside the boxes) and the spread (interquartile range) were shown for each sample. **C.** The PCA showed some overlap among samples. **D.** Correlation heat map showed correlation coefficients between the samples. Correlations close to 1 indicate strong similarity. Heat map showed some correlation between groups. N=4.

