## Supplementary material for "miR-379-3p counteracts cancer cachexia through regulation of pyrimidinergic receptor, mitochondrial stress and interferon response": Figures and supplementary figures: miR-379 supplementary tables.pdf

| Group | PBS F | LLC F |
|---|---|---|
| Body weight (BW) (g) | 20.7±0.67 | 21.6±0.88 |
| Tumour weight (TW) (g) | N/A | 0.92±0.27 |
| BW - TW (g) | 20.7±0.67 | 20.7±0.62 |
| TA (g) | 0.049±0.0045 | 0.033±0.0021 |
| EDL (g) | 0.0134±0.0052 | 0.010±0.0032 |
| GAS (g) | 0.114±0.0119 | 0.09±0.015 |
| SOL (g) | 0.0062±0.0006 | 0.006±0.0011 |
| QUAD (g) | 0.15±0.0083 | 0.14±0.0139 |

Supplementary Table 1.

**Changes in muscle and body size in female mice three weeks after injection of PBS/LLC cells initiating tumour growth. Data represented as mean ± SEM.** Data represented as mean ± SEM. Average body weight and tumour size represented. Average mass of left and right muscle is represented. TA, tibialis anterior, EDL, extensor digitorum longus, GAS, gastrocnemius, SOL, soleus, QUAD, quadriceps.

| Group | PBS M | LLC M |
|---|---|---|
| Body weight (BW) (g) | 35.6±1.12 | 34.6±1.5 |
| Tumour weight (TW) (g) | N/A | 0.07±0.45 |
| BW - TW (g) | 35.6±1.121 | 34.59±1.52 |
| TA (g) | 0.057±0.002 | 0.05±0.002 |
| EDL (g) | 0.012±0.002 | 0.018±0.001 |
| GAS (g) | 0.16±0.006 | 0.129±0.008 |
| SOL (g) | 0.007±0.0008 | 0.04*±0.015 |
| QUAD (g) | 0.16±0.014 | 0.138±0.02 |

Supplementary Table 2.

**Changes in muscle and body size in male mice three weeks after injection of PBS/LLC cells initiating tumour growth. Data represented as mean ± SEM.** Data represented as mean ± SEM. Average body weight and tumour size represented. Average mass of left and right muscle is represented. TA, tibialis anterior, EDL, extensor digitorum longus, GAS, gastrocnemius, SOL, soleus, QUAD, quadriceps.

**Supplementary Table 3. The overlap between MitoCarta genes and proteins regulated in cachexia on males and females.**

Table shows up- or downregulated proteins (High/Low) in the LLC group as compared to the PBS group, type of sample (M - male or F-female), description of gene function, location within mitochondria and mitochondrial pathway where they are involved. MIM (mitochondrial inner membrane); MOM (mitochondrial outer membrane). IMS (inner mitochondrial space).

| Protein | High/Low | M/F | Description | Mito Localization | Mito Pathways |
|---|---|---|---|---|---|
| PDHB | *Low* | *M* | *pyruvate dehydrogenase (lipoamide) beta* | *Matrix* | *Metabolism > Carbohydrate metabolism > Pyruvate metabolism* |
| UQCR10 | *Low* | *M* | *ubiquinol-cytochrome c reductase, complex III subunit X* | *MIM* | *OXPHOS > Complex III > CIII subunits \| OXPHOS > OXPHOS subunits* |
| NDUFS1 | *Low* | *M* | *NADH:ubiquinone* | *MIM* | *OXPHOS > Complex* |

| | | | | | |
|---|---|---|---|---|---|
| | | | *oxidoreductase core subunit S1* | | *I > CI subunits \| Metabolism > Metals and cofactors > Fe-S-containing proteins \| OXPHOS > OXPHOS subunits* |
| PMPCA | *Low* | *M* | *peptidase (mitochondrial processing) alpha* | *Matrix* | *Protein import, sorting and homeostasis > Protein import and sorting > Preprotein cleavage \| Protein import, sorting and homeostasis > Protein homeostasis > Proteases* |
| RTN4IP1 | *Low* | *M* | *reticulon 4 interacting protein 1* | *Matrix* | *N/A* |
| NDUFS2 | *Low* | *M* | *NADH:ubiquinone oxidoreductase core subunit S2* | *MIM* | *OXPHOS > Complex I > CI subunits \| Metabolism > Metals and cofactors > Fe-S-containing proteins \| OXPHOS > OXPHOS subunits* |
| NDUFV1 | *Low* | *M* | *NADH:ubiquinone oxidoreductase core subunit V1* | *MIM* | *OXPHOS > Complex I > CI subunits \| Metabolism > Metals and cofactors > Fe-S-containing proteins \| OXPHOS > OXPHOS subunits* |
| PRDX3 | *Low* | *M* | *peroxiredoxin 3* | *Matrix* | *Metabolism > Detoxification > ROS and glutathione metabolism* |
| TIMM44 | *Low* | *M* | *translocase of inner mitochondrial membrane 44* | *MIM* | *Protein import, sorting and homeostasis > Protein import and sorting > Import motor* |
| MTCH2 | *Low* | *M* | *mitochondrial carrier 2* | *MOM* | *Small molecule transport > SLC25A family \| Mitochondrial dynamics and surveillance > Fusion* |
| VDAC1 | *Low* | *M* | *voltage-dependent anion channel 1* | *MOM* | *Signaling > Calcium homeostasis > Mitochondrial permeability* |

| | | | | | |
|---|---|---|---|---|---|
| | | | | | *transition pore \| Mitochondrial dynamics and surveillance > Organelle contact sites \| Small molecule transport* |
| NDUFA2 | *Low* | *M* | *NADH:ubiquinone oxidoreductase subunit A2* | *MIM* | *OXPHOS > Complex I > CI subunits \| OXPHOS > OXPHOS subunits* |
| COX4I1 | *Low* | *M* | *cytochrome c oxidase subunit 4I1* | *MIM* | *OXPHOS > Complex IV > CIV subunits \| OXPHOS > OXPHOS subunits* |
| IVD | *Low* | *M* | *isovaleryl coenzyme A dehydrogenase* | *Matrix* | *Metabolism > Amino acid metabolism > Branched-chain amino acid metabolism* |
| COQ8A | *Low* | *M* | *coenzyme Q8A* | *MIM* | *Metabolism > Metals and cofactors > Coenzyme Q metabolism* |
| NDUFA10 | *Low* | *M* | *NADH:ubiquinone oxidoreductase subunit A10* | *MIM* | *OXPHOS > Complex I > CI subunits \| OXPHOS > OXPHOS subunits* |
| CLPP | *Low* | *M* | *caseinolytic mitochondrial matrix peptidase proteolytic subunit* | *Matrix* | *Protein import, sorting and homeostasis > Protein homeostasis > Proteases* |
| HSD17B10 | *Low* | *M* | *hydroxysteroid (17-beta) dehydrogenase 10* | *Matrix* | *Mitochondrial central dogma > mtRNA metabolism > mtRNA granules \| Mitochondrial central dogma > mtRNA metabolism > Polycistronic mtRNA processing \| Mitochondrial central dogma > mtRNA metabolism > mt-tRNA modifications \| Metabolism > Lipid metabolism > Fatty acid oxidation \| Metabolism > Lipid metabolism >* |

| | | | | | |
|---|---|---|---|---|---|
| | | | | | *Cholesterol, bile acid, steroid synthesis \| Metabolism > Amino acid metabolism > Branched-chain amino acid metabolism* |
| TUFM | *Low* | *M* | *Tu translation elongation factor, mitochondrial* | *Matrix* | *Mitochondrial central dogma > Translation > Translation factors* |
| DECR1 | *Low* | *M* | *2,4-dienoyl CoA reductase 1, mitochondrial* | *Matrix* | *Metabolism > Lipid metabolism > Fatty acid oxidation* |
| NDUFA13 | *Low* | *M* | *NADH:ubiquinone oxidoreductase subunit A13* | *MIM* | *OXPHOS > Complex I > CI subunits \| OXPHOS > OXPHOS subunits* |
| ATP5MPL | *Low* | *M* | *ATP synthase membrane subunit 6.8PL* | *MIM* | *OXPHOS > Complex V > CV subunits \| OXPHOS > OXPHOS subunits* |
| HADHB | *Low* | *M* | *hydroxyacyl-CoA dehydrogenase trifunctional multienzyme complex subunit beta* | *MIM* | *Metabolism > Carbohydrate metabolism > Ketone metabolism \| Metabolism > Lipid metabolism > Fatty acid oxidation \| Metabolism > Amino acid metabolism > Lysine metabolism* |
| GRHPR | *Low* | *M* | *glyoxylate reductase/hydroxypyruvate reductase* | *Matrix* | *Metabolism > Amino acid metabolism > Glyoxylate metabolism* |
| CYCS | *High* | *M* | *cytochrome c, somatic* | *IMS* | *Metabolism > Metals and cofactors > Heme-containing proteins \| Metabolism > Electron carriers > Cytochromes \| OXPHOS > OXPHOS subunits \| OXPHOS > Cytochrome C \| Mitochondrial dynamics and surveillance > Apoptosis* |
| DBI | *High* | *M* | *diazepam binding inhibitor* | *unknown* | *Metabolism > Lipid metabolism* |

| | | | | | |
|---|---|---|---|---|---|
| HINT1 | *High* | *M* | *histidine triad nucleotide binding protein 1* | *unknown* | *Metabolism > Nucleotide metabolism > Nucleotide synthesis and processing* |
| PMPCA | *Low* | *F* | *peptidase (mitochondrial processing) alpha* | *Matrix* | *Protein import, sorting and homeostasis > Protein import and sorting > Preprotein cleavage \| Protein import, sorting and homeostasis > Protein homeostasis > Proteases* |
| SLC25A20 | *Low* | *F* | *solute carrier family 25 (mitochondrial carnitine/acylcarnitine translocase), member 20* | *MIM* | *Metabolism > Lipid metabolism > Fatty acid oxidation \| Metabolism > Metals and cofactors > Carnitine synthesis and transport \| Metabolism > Metals and cofactors > Carnitine shuttle \| Small molecule transport > SLC25A family* |
| PHB2 | *Low* | *F* | *prohibitin 2* | *MIM* | *Protein import, sorting and homeostasis > Protein homeostasis > Chaperones* |
| CHCHD3 | *Low* | *F* | *coiled-coil-helix-coiled-coil-helix domain containing 3* | *MIM* | *Mitochondrial dynamics and surveillance > Cristae formation > MICOS complex \| Mitochondrial dynamics and surveillance > Intramitochondrial membrane interactions* |
| SLC25A5 | *Low* | *F* | *solute carrier family 25 (mitochondrial carrier, adenine nucleotide translocator), member 5* | *MIM* | *Metabolism > Nucleotide metabolism > Nucleotide import \| Signaling > Calcium homeostasis > Mitochondrial permeability* |

| | | | | | |
|---|---|---|---|---|---|
| | | | | | *transition pore \| Small molecule transport > SLC25A family* |
| GRSF1 | *Low* | *F* | *G-rich RNA sequence binding factor 1* | *Matrix* | *Mitochondrial central dogma > mtRNA metabolism > mtRNA granules \| Mitochondrial central dogma > mtRNA metabolism > Polycistronic mtRNA processing \| Mitochondrial central dogma > mtRNA metabolism > mtRNA stability and decay \| Mitochondrial central dogma > Translation > Mitochondrial ribosome assembly* |
| PPIF | *High* | *F* | *peptidylprolyl isomerase F (cyclophilin F)* | *Matrix* | *Signaling > Calcium homeostasis > Mitochondrial permeability transition pore* |
| HINT2 | *High* | *F* | *histidine triad nucleotide binding protein 2* | *Matrix* | *Metabolism > Lipid metabolism > Cholesterol, bile acid, steroid synthesis* |
| SYNJ2BP | *High* | *F* | *synaptojanin 2 binding protein* | *MOM* | *Mitochondrial dynamics and surveillance > Organelle contact sites* |

**Supplementary Table 4. Mitochondrial proteins (MitoCarta) regulated by miR-39-3p in cachectic mice.** Table presents the overlap between proteins regulated in male (M) and female (F) LLC miR -379-3p-treated mice and transcripts regulated in muscle from female LLC miR-379-3p-treated mice. Table shows protein or gene name, up or downregulation (Up/down) in proteomics (Prot) or RNA-seq data (Gene), and type of sample (M or F), description of gene function, location within mitochondria and mitochondrial pathway where they are involved. MIM (mitochondrial inner membrane); MOM (mitochondrial outer membrane). IMS (inner mitochondrial space).

| *Gene* | *Up/ Down* | *M/ F* | *Prot/ Gene* | *Description* | *Mito Localisation* | *Mito Pathways* |
|---|---|---|---|---|---|---|
| *Uqcrc1* | Up | M | Prot | Ubiquinol-cytochrome c reductase core protein 1 | MIM | Protein import, sorting and homeostasis > Protein import and |

| | | | | | | |
|---|---|---|---|---|---|---|
| | | | | | | sorting > Preprotein cleavage \| OXPHOS > Complex III > CIII subunits \| OXPHOS > OXPHOS subunits |
| *Coq5* | Up | M | Prot | Coenzyme Q5 methyltransferase | MIM | Metabolism > Metals and cofactors > Coenzyme Q metabolism |
| *Uqcrq* | Up | M | Prot | Ubiquinol-cytochrome c reductase, complex III subunit VII | MIM | OXPHOS > Complex III > CIII subunits \| OXPHOS > OXPHOS subunits |
| *Dbt* | Up | M | Prot | Dihydrolipoamide branched chain transacylase E2 | Matrix | Metabolism > Amino acid metabolism > Branched-chain amino acid metabolism \| Metabolism > Amino acid metabolism > Branched-chain amino acid dehydrogenase complex |
| *Dlat* | Up | M | Prot | Dihydrolipoamide S-acetyltransferase (E2 component of pyruvate dehydrogenase complex) | Matrix | Metabolism > Carbohydrate metabolism > Pyruvate metabolism |
| *Pdhx* | Up | M | Prot | Pyruvate dehydrogenase complex, component X | Matrix | Metabolism > Carbohydrate metabolism > Pyruvate metabolism |
| *Afg3l2* | Up | M | Prot | AFG3-like AAA ATPase 2 | MIM | Protein import, sorting and homeostasis > Protein homeostasis > Proteases |
| *Echs1* | Up | M | Prot | Enoyl Coenzyme A hydratase, short chain, 1, mitochondrial | Matrix | Metabolism > Lipid metabolism > Fatty acid oxidation \| Metabolism > Amino acid metabolism > Branched-chain amino acid metabolism \| Metabolism > Amino acid metabolism > Lysine metabolism |
| *Abhd11* | Up | M | Prot | Abhydrolase domain containing 11 | Matrix | Metabolism > Carbohydrate metabolism > TCA- |

| | | | | | | |
|---|---|---|---|---|---|---|
| | | | | | | associated |
| *Ndufb9* | Up | M | Prot | NADH:ubiquinone oxidoreductase subunit B9 | MIM | OXPHOS > Complex I > CI subunits \| OXPHOS > OXPHOS subunits |
| *Timm44* | Up | M | Prot | Translocase of inner mitochondrial membrane 44 | MIM | Protein import, sorting and homeostasis > Protein import and sorting > Import motor |
| *Ndufa2* | Up | M | Prot | NADH:ubiquinone oxidoreductase subunit A2 | MIM | OXPHOS > Complex I > CI subunits \| OXPHOS > OXPHOS subunits |
| *Timm8b* | Up | M | Prot | Translocase of inner mitochondrial membrane 8B | MIM | Protein import, sorting and homeostasis > Protein homeostasis > Chaperones \| Protein import, sorting and homeostasis > Protein import and sorting |
| *Acat1* | Up | M | Prot | Acetyl-Coenzyme A acetyltransferase 1 | Matrix | Metabolism > Carbohydrate metabolism > Ketone metabolism \| Metabolism > Lipid metabolism > Fatty acid oxidation \| Metabolism > Amino acid metabolism > Branched-chain amino acid metabolism \| Metabolism > Amino acid metabolism > Lysine metabolism |
| *Mrpl49* | Up | M | Prot | Mitochondrial ribosomal protein L49 | Matrix | Mitochondrial central dogma > Translation > Mitochondrial ribosome |
| *Ndufb10* | Up | M | Prot | NADH:ubiquinone oxidoreductase subunit B10 | MIM | OXPHOS > Complex I > CI subunits \| OXPHOS > OXPHOS subunits |
| *Coq8a* | Up | M | Prot | Coenzyme Q8A | MIM | Metabolism > Metals and cofactors > Coenzyme Q metabolism |
| *Aldh6a1* | Up | M | Prot | Aldehyde | MIM | Metabolism > |

| | | | | | | |
|---|---|---|---|---|---|---|
| | | | | dehydrogenase family 6, subfamily A1 | | Amino acid metabolism > Branched-chain amino acid metabolism |
| *Ppif* | Up | M | Prot | Peptidylprolyl isomerase F (cyclophilin F) | Matrix | Signaling > Calcium homeostasis > Mitochondrial permeability transition pore |
| *Ech1* | Up | M | Prot | Eoyl coenzyme A hydratase 1, peroxisomal | Matrix | Metabolism > Lipid metabolism |
| *Idh2* | Up | M | Prot | Isocitrate dehydrogenase 2 (NADP+), mitochondrial | Matrix | Metabolism > Carbohydrate metabolism > TCA cycle |
| *Aldh2* | Up | M | Prot | Aldehyde dehydrogenase 2, mitochondrial | Matrix | Metabolism > Detoxification > Xenobiotic metabolism |
| *Tufm* | Up | M | Prot | Tu translation elongation factor, mitochondrial | Matrix | Mitochondrial central dogma > Translation > Translation factors |
| *Dhrs4* | Up | M | Prot | Dehydrogenase/reductase (SDR family) member 4 | Matrix | Metabolism > Vitamin metabolism > Vitamin A metabolism |
| *Ndufa12* | Up | M | Prot | NADH:ubiquinone oxidoreductase subunit A12 | MIM | OXPHOS > Complex I > CI subunits \| OXPHOS > OXPHOS subunits |
| *Hadhb* | Up | M | Prot | Hydroxy acyl-CoA dehydrogenase trifunctional multienzyme complex subunit beta | MIM | Metabolism > Carbohydrate metabolism > Ketone metabolism \| Metabolism > Lipid metabolism > Fatty acid oxidation \| Metabolism > Amino acid metabolism > Lysine metabolism |
| *Mpst* | Up | M | Prot | Mercaptopyruvate sulfurtransferase | MIM | Metabolism > Detoxification > Xenobiotic metabolism \| Metabolism > Sulfur metabolism |
| *Slc25a4* | Down | M | Prot | Solute carrier family 25 (mitochondrial carrier, adenine nucleotide translocator), member 4 | MIM | Metabolism > Nucleotide metabolism > Nucleotide import \| Signaling > Calcium homeostasis > |

| | | | | | | |
|---|---|---|---|---|---|---|
| | | | | | | Mitochondrial permeability transition pore \| Small molecule transport > SLC25A family |
| *Pccb* | Down | M | Prot | Propionyl Coenzyme A carboxylase, beta polypeptide | Matrix | Metabolism > Carbohydrate metabolism > Propanoate metabolism \| Metabolism > Lipid metabolism > Fatty acid oxidation |
| *Acad10* | Down | M | Prot | Acyl-Coenzyme A dehydrogenase family, member 10 | Matrix | Metabolism > Lipid metabolism > Fatty acid oxidation |
| *Cbr2* | Down | M | Prot | Carbonyl reductase 2 | Matrix | Metabolism |
| *Ahcyl1* | Down | M | Prot | S-adenosylhomocysteine hydrolase-like 1 | MOM | Mitochondrial dynamics and surveillance > Organelle contact sites |
| *Lrpprc* | Up | F | Prot | Leucine-rich PPR-motif containing | Matrix | Mitochondrial central dogma > mtRNA metabolism > mtRNA stability and decay \| Mitochondrial central dogma > Translation |
| *Ethe1* | Up | F | Prot | Ethylmalonic encephalopathy 1 | Matrix | Metabolism > Sulfur metabolism |
| *Glud1* | Up | F | Prot | Glutamate dehydrogenase 1 | Matrix | Metabolism > Amino acid metabolism > Glutamate metabolism \| Metabolism > Amino acid metabolism > GABA metabolism |
| *Ptges2* | Up | F | Prot | Prostaglandin E synthase 2 | MIM | Metabolism > Lipid metabolism > Eicosanoid metabolism |
| *Txnrd2* | Up | F | Prot | Thioredoxin reductase 2 | Matrix | Metabolism > Detoxification > ROS and glutathione metabolism \| Metabolism > Detoxification > Selenoproteins |
| *Nit2* | Up | F | Prot | Nitrilase family, member 2 | unknown | Metabolism > Detoxification |
| *Fam210a* | Up | F | Prot | Family with sequence similarity 210, member A | MIM | N/A |
| *Gsr* | Up | F | Prot | Glutathione reductase | Matrix | Metabolism > |

| | | | | | | |
|---|---|---|---|---|---|---|
| | | | | | | Detoxification > ROS and glutathione metabolism |
| *Fam162a* | Up | F | Prot | Family with sequence similarity 162, member A | MIM | N/A |
| *Acp6* | Up | F | Prot | Acid phosphatase 6, lysophosphatidic | IMS | Metabolism > Lipid metabolism > Phospholipid metabolism |
| *Cox6a2* | Down | F | Prot | Cytochrome c oxidase subunit 6A2 | MIM | OXPHOS > Complex IV > CIV subunits \| OXPHOS > OXPHOS subunits |
| *Pdp1* | Down | F | Prot | Pyruvate dehyrogenase phosphatase catalytic subunit 1 | MIM | Metabolism > Carbohydrate metabolism > Pyruvate metabolism |
| *Slc25a25* | Up | F | Gene | Solute carrier family 25 (mitochondrial carrier, phosphate carrier), member 25 | MIM | Metabolism > Nucleotide metabolism > Nucleotide import \| Signalling > Calcium homeostasis > EF hand proteins \| Small molecule transport > SLC25A family |
| *mt-Atp8* | Up | F | Gene | ATP synthase F0 subunit 8 | MIM | OXPHOS > Complex V > CV subunits \| OXPHOS > OXPHOS subunits |
| *Slc25a10* | Down | F | Gene | Solute carrier family 25 (mitochondrial carrier, dicarboxylate transporter), member 10 | MIM | Metabolism > Carbohydrate metabolism > Gluconeogenesis \| Metabolism > Carbohydrate metabolism > Malate-aspartate shuttle \| Small molecule transport > SLC25A family |
| *Acsm1* | Down | F | Gene | Acyl-CoA synthetase medium-chain family member 1 | Matrix | Metabolism > Lipid metabolism > Fatty acid oxidation |

### Table S5. Sequences

| | | |
|---|---|---|
| Primers Genes | | |
| mB2M F | Sigma | 5’ – GGA GAA TGG GAA GCC GAA CA – 3’ |

| | | |
|---|---|---|
| mB2M R | Sigma | 5’ – TCT CGA TCC CAG TAG ACG GT – 3’ |
| mBak1 F | Sigma | 5'-CCCAACAGCATCTTGGGTCA– 3’ |
| mBak1 R | Sigma | 5'-TGGAACTCTGTGTCGTAGCG– 3’ |
| mBax F | Sigma | 5'-GAACCATCATGGGCTGGACA– 3’ |
| mBax R | Sigma | 5'-AGCCACCCTGGTCTTGGAT– 3’ |
| mBcl2 F | Sigma | 5'-ACTTCTCTCGTCGCTACCGT– 3’ |
| mBcl2 R | Sigma | 5'-TCATTCAACCAGACATGCACCT– 3’ |
| mCoxI F | Sigma | 5’ – CAC TAA TAA TCG GAG CCC CA – 3’ |
| mCoxI R | Sigma | 5’ – TTC ATC CTG TTC CTG CTC CT – 3’ |
| mifi44 F | Sigma | 5’-CTGATTACAAAAGAAGACATGACAGAC-3’ |
| mifi44 R | Sigma | 5’-AGGCAAAACCAAAGACTCCA-3’ |
| mifit1 F | Sigma | 5’-CAAGGCAGGTTTCTGAGGAG-3’ |
| mifit1 R | Sigma | 5’-GACCTGGTCACCATCAGCAT-3’ |
| mIL-18 F | Sigma | 5'-TCAGACAACTTTGGCCGACT– 3’ |
| mIL-18 R | Sigma | 5'-GGTGGATCCATTTCCACTTTGA– 3’ |
| mIL-1B F | Sigma | 5'-ATGCCACCTTTTGACAGTGATG– 3’ |
| mIL-1B R | Sigma | 5'-AAGGTCCACGGGAAAGACAC– 3’ |
| mIP3R1 F | Sigma | 5'-AAGCGGATGGACCTGGTGTTAGAACTG–3’ |
| mIP3R1 R | Sigma | 5'-AATTTGTGCTGTGTGCTTCGCGTAGAACT–3’ |
| mLc3b F | Sigma | 5’ – CAT GCC GTC CGA GAA GAC CT – 3’ |
| mLc3b R | Sigma | 5’ – CGC TCT ATA ATC ACT GGG ATC TT GG – 3’ |
| mNd 1 R | Sigma | 5’ – GAG GCT GTT GCT TGT GTG AC – 3’ |
| mNd1 F | Sigma | 5’ – TCT CGA TCC CAG TAG ACG GT – 3’ |

| | | |
|---|---|---|
| mNLRP3 F | Sigma | 5'-CAGAAGGAAGTGGACTGCGA– 3’ |
| mNLRP3 R | Sigma | 5'-ATGTACAGTCACAGTTTCTGGAG– 3’ |
| mP2RY6 F | Sigma | 5'-GGTAGCGCTGGAAGCTAATG– 3’ |
| mP2RY6 R | Sigma | 5'- TTTCAAGCGACTGCTGCTAA– 3’ |
| mP62 F | Sigma | 5’ – GAG GCA CCC CGA AAC ATGG – 3’ |
| mP62 R | Sigma | 5’-TTC CAC CAA GAG CAA GTAT-3’ |
| mPGC1α F | Sigma | 5’ –TTC CAC CAA GAG CAA GTAT – 3’ |
| mPGC1α R | Sigma | 5’ –CGC TGT CCC ATG AGG TATT – 3’ |
| mRyr1 F | Sigma | 5'-ACCCCACATGGGTTTGAGAC– 3’ |
| mRyr1 R | Sigma | 5'-GACTCCTGACCAGTGTGCTC– 3’ |
| mS29 F | Sigma | 5’ – ATGGGTCACCAGCAGCTCTA– 3’ |
| mS29 R | Sigma | 5’ –GTATTTGCGGATCAGACCGT– 3’ |
| mSERCA2A F | Sigma | 5'-GGCCCGAAACTACCTGGAGC– 3’ |
| mSERCA2A R | Sigma | 5'-CAACGCACATGCACGCACCC– 3’ |
| mTLR-4 F | Sigma | 5'-TGACACCGGGAAGCTTGAAT– 3’ |
| mTLR-4 R | Sigma | 5'-TGTCATCAGGGACTTTGCTGAG– 3’ |
| mTNF- F | Sigma | 5’-GTCAGTCATCTTCTCGAAC-3’ |
| mTNF- R | Sigma | 5’-CAGATAGATGGGCTCATAC-3’ |
| mTom20 R | Sigma | 5’ – GCC TTT TGC GGT CGA AGT AG – 3’ |
| mTom20 F | Sigma | 5’ – AGT CGA GCG AAG ATG GTGG – 3’ |
| miR Primers | | |
| miR-379-3p miRCURY primer | Qiagen | YP00204345 |
| Snord68 miRCURY primer | Qiagen | YP00203911 |

| | | |
|---|---|---|
| Unisp6 miRCURY primer | Qiagen | YP00203954 |
| miR/Scr sequences | | |
| Scrambled (Scr) control | Horizon | 5’- CUCGUUCCUGGUCGUCACCAGU-3’ |
| miR-379-3p mimics | Horizon | 5'-UAUGUAACAUGGUCCACUAACU-3' |

### Table S6. Antibodies and reagents

| Antibodies/Cell probes/Staining | | |
|---|---|---|
| 4′,6-diamidino-2-phenylindole (DAPI) | Sigma | 268298 |
| Alexa Fluor 488nM (highly sensitive green-fluorescent dye) | Invitrogen | A21042 |
| Alexa Fluor for myosin type MYH4 MHc-IIb | Invitrogen | A-21042 |
| Alexa Fluor for myosin type MYH7 MHC-I | Invitrogen | A-21144 |
| Alexa Fluor for myosin type: MYH2 MHC-IIa | Invitrogen | A-21240 |
| Anti-BNIP3 | Abcam | ab109362 |
| Anti-CASPASE3 | Abcam | ab32042 |
| Anti-LAMP1 | Cell Signaling Technology | 9091 |
| Anti-LC3II/I | Abcam | ab10912 |
| Anti-MF20 | DSHB | MF20-C |
| Anti-MFN2 | Cell Signaling Technology | 9482 |
| Anti-OPA1 | Cell Signaling Technology | 80471 |
| Anti-P62 | Cell Signaling | 3.00E+11 |
| Anti-PARKIN | Abcam | ab15954 |
| Anti-TOM20 | Cell Signaling Technology | 42406 |
| ATP6V0A1 | Abnova | H00000535A01 |

| | | |
|---|---|---|
| NFKB | Abcam | AB32360 |
| mTOR | Cell signalling | 2983 |
| pMTOR | Cell signalling | 5536 |
| AMPK | Abcam | ab32047 |
| pAMPK | Cell signalling | 2531 |
| TFEB | abcam | Ab264421 |
| cGAS | Cell signalling | 31659 |
| Hoechst | | 33342 |
| IRDye 800CW Goat anti-Mouse IgG | LI-COR Biosciences | 925-32210 |
| IRDye 800CW Goat anti-Rabbit IgG | LI-COR Biosciences | 926-32211 |
| MHC-I | DHSB | BA-D5 |
| MHC-IIa | DHSB | SC-71 |
| MHc-IIb | DHSB | BF-F3 |
| MitoSOX™ | Invitrogen | M7512 |
| MitoTrackerTM Green FM | Invitrogen | M7514 |
| MitoTrackerTM Red CMXRos | Invitrogen | M7512 |
| MitoTrackerTM Red FM | Invitrogen | M22425 |
| TMRM | Invitrogen | T668 |

### Table S7. Reagents

| Reagent | Manufacturer | Reference |
|---|---|---|
| Experimental Models: Cell Lines | | |
| C2C12 | ATCC | CRL-1772 |
| LLCs | ATCC | CRL-1642 |

| | | |
|---|---|---|
| Experimental Models | | |
| C57BL/6J | Charles River | N/A |
| Materials/Reagents | | |
| 2-Propanol | Merck | I9516 |
| 25X dNTP Mix (100 mM) | Thermo Fisher | 4368814 |
| 3PRIMEG/02 48 x 0.2ml, 3 PrimeG Gradient Thermal Cycler | Techne | 93945-09 |
| Acetic acid | Sigma | A6283 |
| Acrylamide | Sigma | A3699 |
| Ammonium Persulfate | Merck | A3678 |
| BeatBox® | PreOmics | N/A |
| Beta-mercaptoethanol | Sigma | M3148 |
| Bovine Serum Albumin BSA | Sigma | A2153 |
| Bromophenol blue | Sigma | B0126 |
| Cell Counting Kit-8 | Sigma | 96992 |
| Cell culture flask, T-175 | Sarstedt | 83.3912 |
| Cell culture flask, T-75 | Sarstedt | 83.3911 |
| Cell culture plate, 12 well | Sarstedt | 83.3921 |
| Chloroform | Sigma | 25666-100ML |
| Dimethyl Sulfoxide | Sigma | 41639 |
| DMEM | Sigma | D6429 |
| Dulbecco's Phosphate Buffered Saline | Sigma | D8537 |
| EDTA | Sigma | ED2SS |
| Electric Pestle | VWR | SCERSP749540-0000 |

| | | |
|---|---|---|
| Epredia 125 ML OCT embedding cryo-embedding Matrix | Epredia | 12678646 |
| Ethanol | Sigma | E7023 |
| EVOS M7000 Imaging System | Thermo Fisher | AMF7000 |
| Fast SYBR™ Green Master Mix | Applied | 4385617 |
| FBS | Sigma | F7524 |
| Film, free of DNase/RNase, material: PO, transparent, optimised for qPCR | Sarstedt | 95.1994 |
| Fisher BioReagents™ EZ-Run™ Pre-stained Rec Protein Ladder | Fisher Bioreagents | BP36031 |
| Glycerol | Sigma | G6279 |
| Glycine | Sigma | G8898 |
| GSM Grip-Strength Meter for mice and rats | Ugo Basile | 47200UB |
| HS | Sigma | H1270 |
| Hydromount | National diagnostics | HS-106 |
| Isopropanol | Sigma | I9516 |
| Laminin | Sigma | L2020 |
| Leica CM3050 S Cryostat | Leica Biosystems | CM3050s |
| M.O.M mouse on mouse blocking | Invitrogen | R37621 |
| Methanol | Sigma | 34860 |
| miR-181a-5p mimics | Horizon | 5'- AACAUUCAACGCUGUCGGUGAGU-3' |
| miR-24-3p Inhibitor (AM-24) | Horizon | 5'- UGGCUCAGUUCAGCAGGAACAG-3' |
| miR-26a-5p mimics | Horizon | 5'- UUCAAGUAAUCCAGGAUAGGCU-3' |
| miR-379-3p mimics | Horizon | 5'-UAUGUAACAUGGUCCACUAACU-3' |
| miRcury LNA RT kit | Qiagen | 339340 |
| miRcury LNA SYBR Green PCR Kit (4000) RT-qPCR kit | Qiagen | 339347 |

| | | |
|---|---|---|
| miRvana™ miRNA Isolation Kit, with Phenol | Invitrogen | AM1561 |
| N,N,N′,N′-Tetramethylethylenediamine | Merck | T9281 |
| NaCl | Sigma | S9888 |
| Nanodrop 2000 | Invitrogen | N/A |
| Odyssey® Fc Imaging System | LI-COR Biosciences | OFC-1025 |
| omniPAGE Mini Vertical Protein Electrophoresis System | Cleaver Scientific | VS10 |
| PCR plate half skirt, 96 well, transparent, Low-Profile | Sarstedt | 72.1981.232 |
| PCR single tube, 0.5 ml, Biosphere® plus | Sarstedt | 72.735.100 |
| Penicillin/Streptomycin | Invitrogen | P4333 |
| Phosphatase Inhibitor | Merck | P0044 |
| PowerEase™ 90W Power Supply (115 VAC) | Fisher Scientific | PS0090 |
| PreOmics iST 96x (96-sample kit) | PreOmics | P.O.00027 |
| Protease Inhibitor Cocktail | Sigma | P8340 |
| Protein Assay Dye Reagent Concentrate | Bio Rad | 5000006 |
| Random Hexamers (50 µM) | Thermo Fisher | N8080127 |
| RiboLock RNase Inhibitor (40U/µL) | Thermo Fisher | EO0381 |
| RIPA Lysis Buffer, 10X | Sigma | 20-188 |
| RNase free water | Sigma | 3098 |
| Scrambled (Scr) control | Horizon | 5’- CUCGUUCCUGGUCGUCACCAGU-3’ |
| Seesaw-Rocker Large 230V | Stuart | 51900-32 |
| Sense Beta Plus Microplate Reader | Hidex | 425-311 |
| Sodium Dodecyl Sulfate | Merck | L3771 |
| StepOnePlus™ Real-Time PCR System | Applied Biosystems | 4376600 |

| | | |
|---|---|---|
| SuperScript™ II Reverse Transcriptase kit (includes 5X FirstStrand Buffer and 0.1 M DTT) | Invitrogen | 18064014 |
| Transfer Pipette | Sarstedt | 861171 |
| Transwell inserts 0.4 mm | Fisher | 10567522 |
| Trizma® base | Sigma | T1503 |
| Trizma® hydrochloride | Merck | T3253 |
| TrypLE™ Express Enzyme | Invitrogen | 12604013 |
| Tween 20 | Merck | P9416 |
| Western blotting membranes, nitrocellulose | Cytiva | GE10600002 |
| Wheat Germ Agglutinin (WGA), Fluorescein | Vector Labs | FL-1021 |
| Software and Algorithms | | |
| DIANA-miRPath v3 and TarBase v7.0 | N/A | (Vlachos et al., 2015) |
| ChEA3_Encode | Mount Sinai Center for Bioinformatics | (Keenan et al., 2019) |
| ENSEMBLE Bio Mart | | (Shannon et al., 2003) |
| GSEA | UCSanDiego | (Subramanian et al., 2005, Mootha et al., 2003) |
| Image Studio Lite | Image Studio Lite | RRID:SCR_013715 |
| ImageJ | NIH | RRID:SCR_003070 |
| MaxQuant (v 1.6.0.16) | N/A | N/A |
| miRDB | N/A | (Chen and Wang, 2020) |
| miRPathDBv2.0 | Universitat Des Saarlandes | (Kehl et al., 2020) |
| NCBI Blast | N/A | N/A |
| PathVisio | Maastricht Centre for Systems Biology (MaCSBio) | N/A |
| Prism 9 | GraphPad Software | (Prism License) |

| | | |
|---|---|---|
| Prostar | N/A | (Wieczorek et al., 2017) |
| R software | N/A | (Team, 2023) |
| R studio | N/A | (Team, 2023) |
| TargetScanMouse8.0 and Human8.0 | N/A | (Agarwal, 2015) |
| UNIPROT | | |
